# SARS-CoV-2 vaccination during pregnancy enhances hippocampal neurogenesis and working memory in offspring via IFN-gamma responsive microglia

**DOI:** 10.1101/2024.07.03.601972

**Authors:** Fangfang Liu, Jiaoling Tang, Juntao Zou, Hao Liu, Zejie Zuo, Lingxiao Wang, Na Wang, Zhihui Li, Ashutosh Kumar, Kaihua Guo, Dandan Hu, Zhibin Yao

## Abstract

In the face of severe adverse outcomes of pandemic Coronavirus disease 2019 (COVID-19) infection, vaccines proved potently immunogenic and safe in humans and are today strongly recommended in pregnancy. This study investigates, in offspring mice, the effect of maternal COVID-19 vaccination on postnatal physical development, behavior and neurogenesis. After inoculation with inactivated COVID-19 vaccine (Vero Cell) at gestational day 14.5, antibodies to the severe acute respiratory syndrome coronavirus 2 (SARS-CoV-2) were detected in serum of both dams and pups. At one month of age, pups born to vaccinated dams, but not the offspring of non-vaccinated dams, exhibited greater working memory and more neural cell proliferation, neuroblast formation, neuronal stem cell activity and larger numbers of mature neurons within the dentate gyrus (DG). Luminex multiplex assay revealed elevated levels of hippocampal cytokines/chemokines critical to neurogenesis and memory function, namely interferon-γ [IFN-γ] and CX3C motif chemokine ligand 1 [CX3CL1]. Conditional knockout technology implicated microglial IFN-γ receptor 1 (IFNγR1) and CX3C motif chemokine receptor 1 (CX3CR1) as crucial intercellular participants in the neuronal developmental process, via regulating microglial activation and chemotaxis, respectively. We propose that, rather than posing risk for neurodevelopmental abnormalities, maternal SARS-CoV-2 vaccination transiently enhances hippocampal neurogenesis and working memory in offspring.

## Introduction

The coronavirus disease 2019 (COVID-19) pandemic caused by the severe acute respiratory syndrome coronavirus 2 (SARS-CoV-2) killed millions of people worldwide^1^. Pregnant women with symptomatic COVID-19 are at increased risk for adverse pregnancy-related complications, preterm births and respiratory distress in their infants ^2–5^. Thus, COVID-19 vaccination has been recommended widely for pregnant women, breastfeeding women, and women contemplating pregnancy.^6^ ^7^. Recommended vaccination regimens vary in different countries. The Centers for Disease Control (CDC) in the US and the American college of obstetricians and gynecologists (ACOG) strongly recommend SARS-CoV-2 vaccination for persons 12 years of age or older, including pregnant women, with an additional booster dose of RNA vaccine when pregnant ^7^. Several studies recently reported that vaccination with SARS-CoV-2 RNA vaccine did not adversely affect pregnancy ^8^ ^9^ or neonatal outcomes ^7^ ^10^ ^11^.

SARS-CoV-2 vaccine was made available to high-risk individuals starting in December 2020, but pregnancy remains a contraindication for SARS-CoV-2 inactivated vaccine in China. Because early clinical trials excluded pregnant participants, little clinical information is available concerning the vaccine’s safety during pregnancy ^12^. Two reports have shown that two-doses of inactivated SARS-CoV-2 vaccine (CoronaVac, Sinovac Life Science) given to female mice before and during pregnancy did not affect the pregnancy or the physical development of pups ^13^ ^14^. Furthermore, maternal immunization had no impact on the results of cognitive function when pups reached adulthood. No in-depth analysis is available concerning hippocampal neurogenesis or neuronal-glial interactions in pups born to pregnancy-vaccinated dams.

We reported earlier that influenza vaccination during pregnancy enhanced neurogenesis and spatial learning abilities in adolescent pups^15^, and improved working memory in pregnant mice^16^. Furthermore, a recent review hypothesized that adult COVID-19 immunization may promote adult hippocampal neurogenesis via microglial activation^17^ ^18^. Brain-resident immune cells continuously react to environmental cues and their release of cytokines IFN-γ, IL-1β, IL-6 or transforming growth factor β (TGF-β) has been implicated in the control of adult hippocampal neurogenesis^19–22^. Neuronal CX3CL1 has a cell-autonomous effect on adult neurogenesis in health, reverses neuronal loss in neurodegenerative disease models^23^ ^24^ and communicates with microglial CX3CR1 in maintaining a homeostatic state^25^ ^26^. Emerging data implicate microglia-neuron interactions in neonatal neuronal development and in adult hippocampal neurogenesis ^27–30^, but underlying mechanisms are unknown.

Here, we present evidence for microglial interaction with immature neurons in mice born of SARS-CoV-2 vaccinated dams and data supporting a role for microglia in neuronal proliferation and differentiation via IFN-γ and CX3CR1 signaling during brain development.

## Materials and methods

### Animals

We used female C57BL/6 mice aged 6-8 weeks and 12-16 weeks and males aged 8-12 weeks Primiparous pregnant mice were obtained by mating the younger females with healthy male mice. 12-16-week-old once-mated mice were impregnated to obtain non-primiparous dams. Detection of a vaginal plug designated gestational day 0 (GD0), and newborn pups were designated postnatal day 0 (P0). We purchased CX3CR1-GFP transgenic (B6.129P2(Cg)-*Cx3cr1*^tm1Litt^/J) and IFN-γR floxed (C57BL/6N-*Ifngr1*^tm1.1Rds^/J) mice from the Jackson Laboratory (Stock #: 005582; 025394). All other mice were purchased from the Laboratory Animal Center of Sun Yat-sen University. Since the coding exon 2 of *CX3CR1* gene was replaced by EGFP, CX3CR1-GFP/GFP mice were deemed lacking CX3CR1. To induce efficient recombination of CreER enzyme and reduce potential side effects, we injected 4-Hydroxytamoxifen (H6278, Sigma-Aldrich, 1 mg/kg i.p., 3 times) into pups at P7 to P9. Microglia depletion was induced pharmacologically by feeding pexidartinib (PLX3397; Absin, #abs810318) formulated diet, containing a CSF1R inhibitor at a concentration of 500 mg/kg diet for 2 weeks ^31^. Dams were housed individually in an IVC cage in a specific pathogen-free (SPF) barrier with 12 h light/dark cycle and food and water provided ad libitum. Pups were weaned at P21 and housed in groups of four to five sex-matched littermates in a SPF barrier room with constant temperature (22 LJ) and relative humidity (40%). To exclude maternal variables, we took one to two pups from individual dams randomly to constitute groups. Animal experiments complied with regulations formulated by the Institutional Animal Ethics Committee of Sun Yat-sen University. Each pregnant dam was kept individually in an IVC cage in an SPF barrier.

### SARS-CoV-2 vaccination

Pregnant mice received on gestational day 14.5 a single intramuscular (quadriceps) injection, 50 μL or 100 μL of inactivated SARS-CoV-2 vaccine (BBIBP-CorV, Sinopharm, Beijing) containing 0.65 U or 1.3 U; controls received an equal volume of sterile phosphate-buffered saline (PBS). We previously reported that the vaccine adjuvant, aluminum hydroxide, does not impact mouse neurogenesis or behavior when administered alone during early life^32^. The inoculation time point was chosen according to maternal immune activation models reported in earlier studies,^32–35^ including our own. Although maternal vaccination is recommended for human subjects at any gestational age, it has been reported that COVID-19 vaccination early in the third trimester elicits higher spike-specific antibody levels in both maternal and umbilical cord blood^36^. The inactivated COVID-19 vaccine (BBIBP-CorV, Sinopharm, Beijing) was produced by China’s national biotechnological group (CNBG, Beijing). The vaccine contains the effective constituent, 19nCov-CDC-Tan-HB02, which protects against SARS-CoV-2. The immunoreactivity and safety of this vaccine have been validated in humans^37^ ^38^.

### SARS-CoV-2 Spike IgM and IgG measurements

Blood samples were collected from dams on day (d) 2, d3, d5, d7, and d14 after vaccination, and from pups at approximately 1 month of age (following behavioral tests at postnatal day 31). Legend MAX^TM^ SARS-CoV-2 Spike RBD IgM human ELISA kit (448307, Bio Legend, San Diego, CA, USA) was used to quantitate the concentration of IgM antibody against SARS-CoV-2 in the serum of pregnant mice and pups. Lacking a commercial mouse ELISA kit, we substituted HRP-conjugated anti-mouse IgM for the anti-human IgM conjugate to detect murine anti-SARS-CoV-2 IgM. The protocol otherwise followed the manufacturer’s instructions. The top IgM standard represented 30 ng/mL; six doubling dilutions were performed using assay buffer, which alone represented 0 ng/mL. The linear relationship (R^2^ value > 98) demonstrated acceptable quality control specifications with the modified protocol. We also assessed the specificity of the serum sample against the SARS-CoV-2 antigen via negative Serum from non-vaccinated pregnant mice and their pups served as negative controls. All mice were tested behaviorally and euthanized for histological analyses when SARS-CoV2 IgM levels in the vaccinated group fell below 5 ng/mL in pups and 20 ng/mL in dams.

Anti-RBD neutralizing antibody ELISA kit (DD3101, Vazyme Biotech, Nanning, China) was used to detect anti-SARS-CoV-2 RBD IgG in mouse sera, starting at 1:10 dilution in assay buffer. The antibody neutralization rate in serum of vaccinated mice was determined spectrophotometrically by plate reader (optical density [OD] 405 nm) using the mean OD450 of negative controls as blank. SARS-CoV-2-neutralizing IgG was deemed positive when the neutralization rate was at least 20%. Individual pregnant mice and pups with undetectable neutralizing IgG in sera were excluded from behavioral studies.

### Body weight

Following SARS-CoV-2 vaccination, the body weight of pregnant mice was recorded each day post-vaccination (E14.5), and the embryo number and birth weights were recorded for the vaccinated and control pup groups.

### Open-field test

As an indicator of anxiety-related behavior changes, ^39^ ^40^ the pups’ exploratory activities were assessed by placing them in an open-field arena of a lighted room to explore for 10 min using TopScanTM 2.0 (Clever Sys., Inc., Reston, VA, USA); video tracking system (Instruments, CA) was used to measure the total distance in the arena and the percentage of time spent in the center. All behavioral tests were performed between hours 10:00 and 17:00 in the light-off phase.

### Novel object recognition

The novel object recognition test included an acquisition trial and a retrieval trial. During the acquisition, one pup was placed in an arena (40 cm × 40 cm × 40 cm blue plastic box) that contained two identical objects (50 mL centrifuge tubes) for 5 min. The exploration was defined as the animal sniffing the object when the distance between its nose and the object was less than 1 cm. The retrieval session was determined 2 h later by replacing one of the objects with a new and distinct object, a Lego piece of different color and shape, reintroducing the mouse to the arena for another, and recording (using TopScanTM 2.0) the time it spent exploring each object in the next 5 min.

### Social preference

We determined social behavior (social approach and social novelty) in three sessions, as we previously described. ^32^ The arena (60 cm long, 40 cm wide, and 22 cm high) was divided into three identical chambers with transparent walls and each side chamber equipped with a clear plexiglass cylinder (7.5 cm diameter) bearing several holes to allow nose contact. During the habitation session, the mouse was allowed to explore freely for 5 min. The mouse was then gently transferred to the middle chamber and the doorways were closed. In the social approach session, the first stranger mouse (stranger 1, C57BL/6 of a different sex) was placed in one of the side chamber cylinders. After opening the doorways, the test mouse was allowed to explore Stranger 1 or the empty cylinder for another 10 min. In the social novelty session, a second stranger C57BL/6 mouse (stranger 2) was placed in the empty cylinder, and the test mouse was allowed to explore the arena, and the two stranger mice for 10 min. Time spent sniffing (which??) the mouse inside of the cylinder was recorded with TopScanTM 2.0.

### Y-maze

Spatial working memory (working memory) ability was assessed in a Y-maze test by analyzing the spontaneous alternation behavior as described in our previous studies ^40^ ^41^. The symmetrical Y-shape apparatus has three arms randomly labeled A, B, and C (respectively, 37 cm long, 6.5 cm wide and 13 cm high). After habituation in the behavioral room, the mouse was placed at the end of an arm(?) and allowed to move freely for 5 min. The activities were recorded with the TopScanTM 2.0 tracking system. Consecutive entries into all three arms (i.e., ABC, ACB, BAC, BCA, CAB, or CBA) were considered an accurate alternation. There, alternation (%) equals the number of correct alternations divided by (the total number of arm entries - 2) 100%. The experimenter and data analyst were blinded to the animal group throughout the test.

### Administration of 5-bromo-2-deoxyuridine (BrdU) and tissue preparation

The protocol was according to the previous research with slight modifications ^5^ ^42^. For neuronal proliferation analysis in the hippocampus, pups were injected once i.p. with BrdU (B9285, Sigma-Aldrich, St. Louis, MO, USA, 50 mg/kg) at P28 and perfused intracardially 24 h later. For neuronal survival and differentiation analysis, another set of pups received four BrdU injections (with a 12 h interval, from P21 to P23) and were euthanized 7 or 28 days after the first BrdU injection. On day 7, the survival of newly immature neurons was analyzed by staining BrdU and DCX. On day 28, matured newborn neurons double labeled with BrdU and NeuN in the dentate gyri (DG) were determined. The mice were anesthetized with 250 mg/kg tribromoethanol (T48402, Sigma-Aldrich) and 2.5% tert-amyl alcohol (240486, Sigma-Aldrich) and were intracardially perfused with pre-cooled 0.9% saline, followed by pre-cooled 4% paraformaldehyde (PFA) 24 h after BrdU injection. Brains were collected, post-fixed overnight in 4% PFA at 4 LJ for 24 h and equilibrated in gradient sucrose solution (15% to 30%) for at least 16 h at 4 LJ. Coronal frozen sections (30 μm) were cut from the hippocampus (SM2000R microtome, Leica Microsystems, Richmond Hill, Ontario, Canada) and stored in cryoprotectant (50% ethylene glycol, 30% sucrose in PBS) at −20 LJ for further histological analysis. For biochemical analyses, hippocampal and cerebral cortical regions of fresh brain tissues were dissected using stereological microscopy, rapidly frozen in liquid nitrogen before storage at −80 LJ. Bilateral hippocampi and cortical tissues from SCV pups and their matched control pups were homogenized in pre-cooled PIPA buffer (Beyotime Biotechnology, 1:5 weight-to-volume ratio) containing inhibitors of proteases and phosphatases. The supernatant collected after centrifugation at 12,000 *g* (4 LJ, 20 min) were frozen prior to analysis.

### Immunofluorescence staining

Brain slices were rinsed in PBS for 5 min, then immersed in citrate antigen retrieval solution (95 ℃, 20 min), incubated in 2 N HCl (30 ℃, 30 min), blocked (37 ℃, 1 h) in 1% donkey serum albumin (DSA) and 0.25% Triton-100 (Sigma-Aldrich, St. Louis, MO, USA), rinsed three times (each 5 min) in 0.1 M boric acid, followed by three PBS washes (5 min). Slices were then incubated (37 ℃, 2 h then overnight, 4 ℃) with primary IgG antibodies: rat anti-Ki67 (14-5698-80, SolA15, 1:400, Invitrogen, Carlsbad, CA, USA), rat anti-BrdU (ab6326, 1: 400; Abcam, Cambridge, MA, USA), goat anti-DCX (sc-8066, 1:400, Santa Cruz Biotechnology, CA, USA), rabbit anti-BDNF (ab108319, 1:200; Abcam, Cambridge, MA, USA), mouse anti-GFAP (G3893, 1:5,000; Sigma-Aldrich, St. Louis, MO, USA), mouse anti-Nestin (MAB353, 1:800; Sigma-Aldrich, St. Louis, MO, USA), mouse anti-Tmem119 (ab209064, 1:400; Abcam, Cambridge, MA, USA), and anti-NeuN (ab104224, 1:1,000; Abcam, Cambridge, MA, USA). The slices were then incubated (37 ℃, 2 h) with species-appropriate secondary antibody: Alexa Fluor 594 donkey anti-rat IgG, Alexa Fluor 488 donkey anti-mouse IgG, Alexa Fluor 488 donkey anti-mouse IgG, or Alexa Fluor 647 donkey anti-rabbit IgG (1:1,000; Invitrogen, Carlsbad, CA, USA).^43^ ^44^. We used ImageJ software (NIH) for fluorescence analysis in the region of interest (dentate gyrus, DG).

### Confocal imaging, 3D reconstruction, and MBF stereoinvestigation

An LSM 780 confocal laser scanning microscope (Zeiss) was used to capture fluorescence immunostaining images using the same parameters to avoid potential technical artifacts. Imaris 3D surface rendering was used according to the method we previously published^45^. Briefly, IBA1, BrdU, and DCX channels were created separately as a surface after automatic contrast adjustment using Imaris software (Bitplane, version 8.4). After 3D reconstruction, a digital zoom tool was used for image magnification for a good display. Microglia and proliferating cells were contacted at less than 0.25 μm from the proliferating cell surface to the microglial surface. Quantitative analyses of Ki67^+^, BrdU^+^DCX^+^, Ki67^+^ DCX^+^, BrdU^+^IBA1^+^, BrdU^+^NeuN^+^, BrdU^+^GFAP^+^, and BrdU^+^Nestin^+^ in the DG were estimated using the optical-fractionator method with unbiased stereology system stereo investigator (Micro Brightfield, Inc., Williston, VT, USA)^15^ ^42^. For tissue sections, a systematic series of 30-μm-thick coronal sections containing the dorsal hippocampus were analyzed (6 brain slices spanning from −1.2 to −2.4 mm in AP axes, from the bregma). The 6 sections were selected from each of the 6 series, with a random start and spanning the whole dorsal hippocampus to ensure that the entire structure was sampled ^46^. A Nikon microscope with a 40 objective was used for all stereological analyses, keeping the coefficient of error (CE) at less than 0.1 to ensure reliable results.

### Multi-plex mouse chemokine/cytokine arrays

We used the Bio-Plex Pro mouse chemokine assay kit (12009159, Bio-Rad), and the manufacturer’s instructions, to determine levels of 31 mouse chemokines/cytokines in mouse blood and homogenized brain tissues. The data were analyzed using the Luminex 200 system (X-200, Bio-Plex).

### Analysis of the scRNA-Seq database

Single-cell RNA sequencing data containing post-vaccination human PBMCs from Arunachalam et al.^47^ was downloaded from the GEO database (accession number GSE171964, (https://www.ncbi.nlm.nih.gov/geo/query/acc.cgi?acc=GSE171964). Seurat (v5) ^48^ was used for the analysis. Briefly, Seurat objects containing human PBMCs collected on day 0, 1, 2, and 7 were created from the feature-barcode matrices and the annotation metadata file provided by the original study. The expression matrices were then normalized and scaled. The principal component analysis (PCA) was conducted, and the top 20 principal components were used in the uniform manifold approximation and projection (UMAP). Differentially expressed genes (DEGs) of each cell cluster were calculated by the *FindMakers* function between different time points. IFN-gamma-related module score was calculated by the *AddModuleScore* function with the input of all genes in the Gene Ontology term Interferon-mediated Signaling Pathway (GO:0140888). GSEA was performed to identify the altered signaling pathways on day 1/day2 after vaccination compared to that on day 0 (Before vaccination). Codes for the analysis are available at https://github.com/LingxiaoWang357/Fangfang_Qi_Vaccine.

## Statistical analysis

The IBM SPSS Statistics 23 software was used for data analysis. GraphPad Prism 8.0 software was used to generate graphs. A two-way repeated ANOVA analysis was used for body weight of both dams and pups. A two-way paired Student’s *t-test* was used to make comparisons for NOR and three-chamber tests. A two-way unpaired Student’s *t-test* was used for two groups. A one-way ANOVA followed by Tukey HSD test was used for three groups. If the data did not meet the criteria of normality by Kolmogorov-Smirnov test, the Mann–Whitney U test was applied. \**p* < 0.05 indicated statistically significant differences.

## Results

### SARS-CoV-2 antibodies were detected in pups born of vaccinated dams

Serum of pregnancy-vaccinated mice had no detectable antibody at 3 days (d) and 7d, regardless of vaccine dose (Fig. 1B and C). However, significant levels of SARS-CoV-2 Spike RBD IgM were detected (at 1:1000 dilution) in serum of vaccinated dams, at approximately 14 d post-vaccination (Figure 1B and C; *P* < 0.01, *P* < 0.01, respectively). The mean levels of viral-specific IgM in high-dose (100 μL) vaccine recipients (31.53 ng/mL) were 8.6-fold greater than the mean serum level in low-dose (50 μL) recipients (301.83 ng/mL). The levels of neutralizing IgG were significantly higher in vaccinated dams and their pups, relative to the controls (neutralization rate > 20%; Figure 1 D-F; *P*<0.01, *P* < 0.001). Thus, the inactivated Sinopharm BBIBP-CorV vaccine elicited strong maternal humoral immune responses, both IgM and neutralizing IgG, and IgG with potential to protect newborn pups crossed the placental barrier within 31 d of maternal vaccination. SARS-CoV-2-specific SCV-IgM antibodies were not detected in pups (data not shown).

**Fig. 1.**
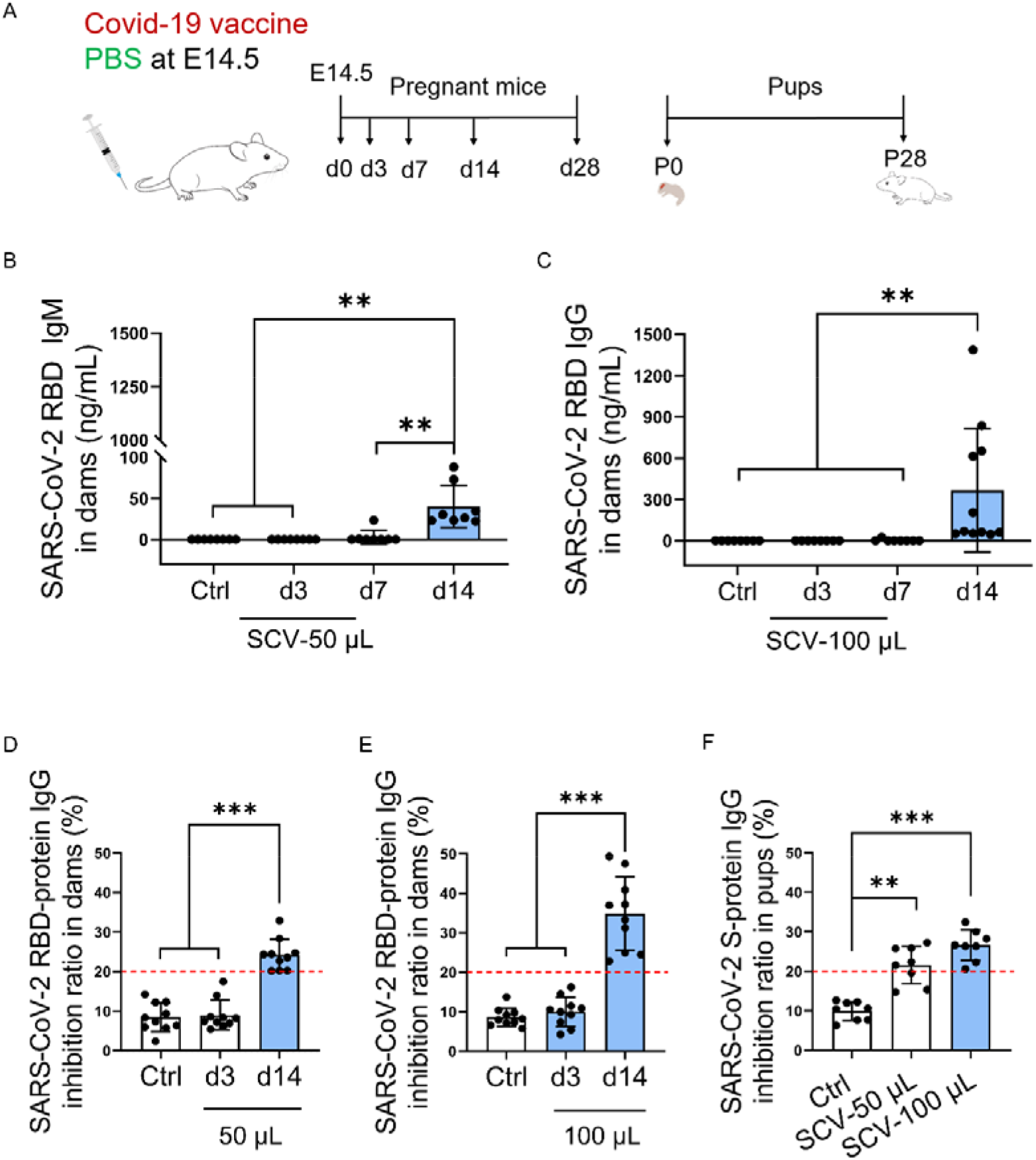
SARS-CoV-2 specific antibodies were detected in both vaccinated pregnant mice and their pups. (A) Diagram of the experimental procedure: pregnant mice were i.m. inoculated with SARS-CoV-2 vaccine with different dosages (50 μL and 100 μL) or PBS at embryonic day 14.5 (E14.5). (B-C) The Spike RBD IgM antibody against SARS-CoV-2 was accurately measured by an ELISA kit in the serum of pregnant mice 3 days, 7 days and 14 days after vaccination, 50 μL for B, 100 μL for C. (D) The S-protein IgG inhibition ratio were assessed by an ELISA kit in the serum of pregnant mice 7 days and 14 days after SARS-CoV2-vaccination 50 (SCV-50) μL for C, 100 (SCV-100) μL for D. (F) The S-protein IgG inhibition ratio were assessed in peripheral serum at postnatal day 31, \*\**p* < 0.01, *\*\*\*p* < 0.001, one-way ANOVA followed by Tukey HSD test; *n*= 8-11 mice/group.

### SARS-CoV-2 vaccination during pregnancy did not affect embryo number nor body weight in pregnant mice or their pups

No significant differences were observed in embryo number (Supplemental Figure 1 A, B, and D) or birth weight among the three groups of neonatal pups (Supplemental Figure 1 E). Likewise, no significant change in the body weights of pregnant mice was attributable to vaccination, regardless of dose (Supplemental Figure 1 C). Although, the body weights of pups born to vaccinated and non-vaccinated dams did not differ significantly in the first 5 postnatal days, from P6 to P25, the low-dose vaccine offspring had significantly greater weight gain than pups of non-vaccinated dams, while weight gain in pups of the high-dose dams was lower than the controls. The body weight differences were not significant for the three groups from postnatal day 26 onward. (Supplemental Figure 1E).

### Offspring of pregnancy-vaccinated dams had increased working memory

Although litter sizes and weight trends did not differ significantly in vaccinated and non-vaccinated pregnant mice, body weight trends in the pups born to vaccinated dams did differ transiently from those born to non-vaccinated dams (Supplemental Figure 1E). Those data encouraged us to investigate whether maternal SARS-CoV2 vaccination might have a discernible effect on the pups’ behavior. Open field testing (OFT) and novel object recognition (NOR tests) were conducted to evaluate the pups’ exploratory behavior and short-term memory function at 1 and 2 months of age. In OFT testing, neither the percentage of time spent in the field center, nor the total distance moved differed significantly in the vaccination and control groups, at either 1 month or 2 months of age, regardless of sex. (Figure 2A-C; Supplemental Figure 2A-C, and Supplemental Figure 3A-C). However, female pups in the vaccination group spent more time than control offspring in the center of the open field arena at age 1 month but not at 2 months (Supplemental Figure 2B). In the NOR task, pups of the vaccinated dams showed a sharp increase in novel object exploration time at 1 month, regardless of sex (Figure 2D-E and Supplemental Figure 2D-E), but not at 2 months (Supplemental Figure 3D-E). In the Y-maze task, at the age of 1 month, the offspring of vaccinated dams spent more time in the new arm than the old arm, as compared with offspring of control dams (Figure. 2F-G, Supplemental Figure 2F-G). No obvious differences were observed in spontaneous alternation, at either 1 month or 2 months (Figure 2I; Supplemental Figure 2F and Supplemental Figure 3F). Similar results were found for total distances moved in the Y maze test and OFT (Figure 2 H and J; Supplemental Figure 2C and G). In the social approach phase, male and female pups born of either vaccinated dams or control dams preferred the stranger mouse to the object at both 1 and 2 months of age (Supplemental Figure 3G and Supplemental Figure 4A, C and D). In the social novelty phase, there was similar preference for the stranger mouse than for the familiar (Supplemental Figure 3H; Supplemental Figure 4 B, E and F). Thus, maternal vaccination did not affect the pups’ social activities, but it did influence short-term memory at age 1 month, regardless of sex.

**Fig. 2.**
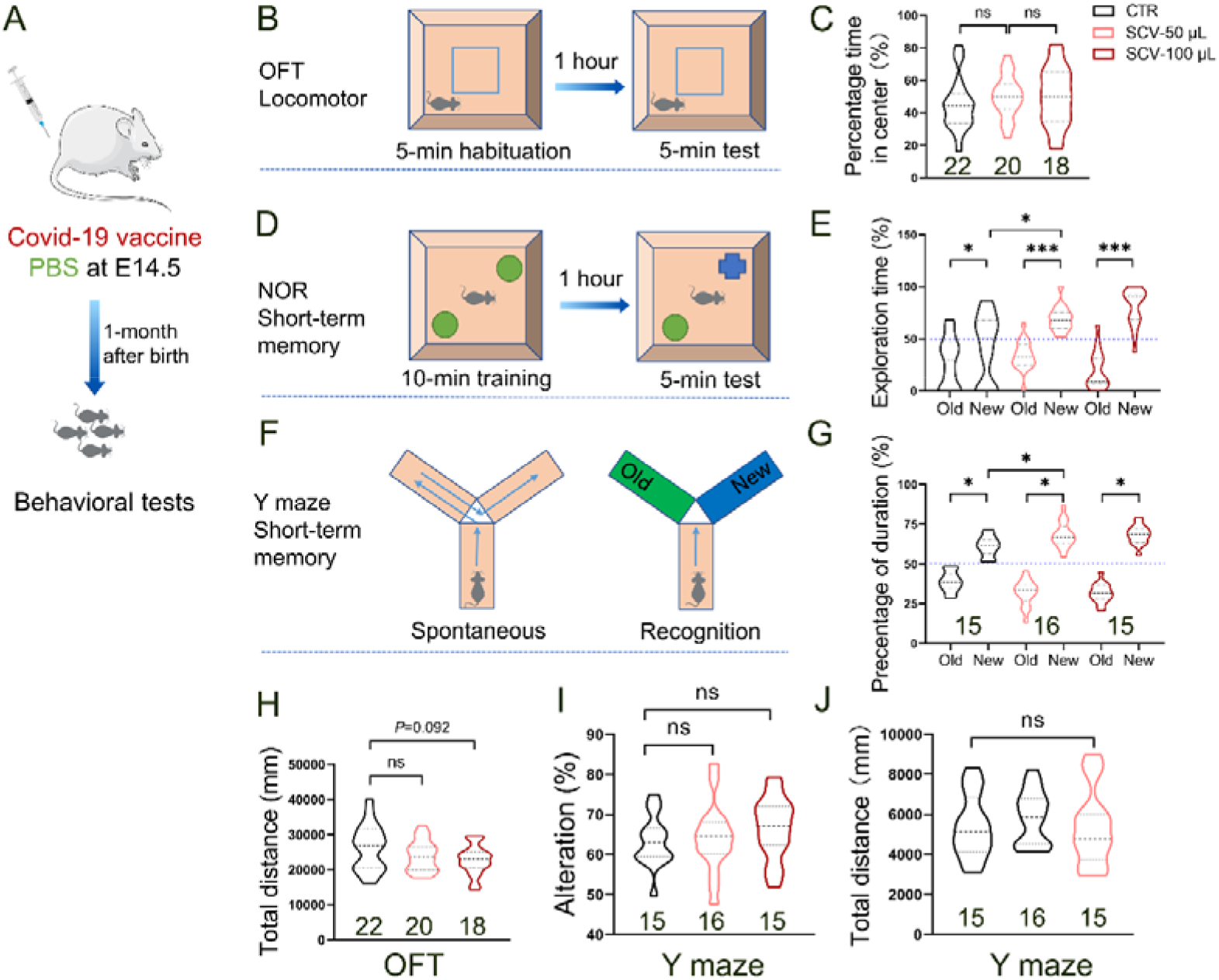
Maternal SARS-CoV-2 vaccination promoted short-memory in pups at 1 month after birth. A Diagram of the experimental procedure, behavior tests were performed 1 month after birth. (B) Experimental approach of habituation and testing for OFT (C) Effects of maternal SARS-CoV-2 vaccination on exploratory behavior in pups (SCV pups) from vaccinated dams and the controls were measured in OFT at 1 month postnatally, \**p* < 0.05 indicate significance by one-way ANOVA followed by Tukey HSD test; (D) Experimental approach of NOR training and testing; (E) At age of 1 month, SARS-CoV-2 vaccination pups showed increased preference to the new object in the NOR tests as compared to vehicle controls; (F) Experimental approach of spontaneous stage and recognition stage in Y maze test; (G) At age of 1 month, SCV pups performed increased duration in novel arm in the Y maze tests compared to the controls; No obvious changes were found in total distances in OFT (H) and Y maze test (J), alteration percentage in spontaneous stage of Y maze test (I) in SCV pups, relative to the controls. Summary data are presented as mean ± s.e.m. ns: non-significant; \*\**p* < 0.01, *\*\*\*p* < 0.001, one-way ANOVA followed by Tukey HSD test; *n*=12 pups/group.

### Hippocampal neurogenesis is increased in pups of SARS-CoV-2 vaccinated dams

Working memory has been linked to neurogenesis in the hippocampus’ dentate gyrus ^49–51^. We therefore investigated the effect of maternal SARS-CoV2 vaccination on neuronal proliferation and differentiation and found that the number of BrdU^+^ neurons in the dentate gyrus of 1 month old pups born of vaccinated dams was significantly greater than in pups of non-vaccinated dams (Figure 3A-B). Double-labeling by the proliferating marker (BrdU^+^) and the neuroblast marker (DCX^+^) allowed us to identify immature neurons amongst newly generated hippocampal cells. We observed that the numbers of newly born neurons (BrdU^+^DCX^+^) and neural stem cells (BrdU^+^Nestin^+^) in the dentate gyrus of pups born of pregnancy-vaccinated dams were significantly greater than in control pups of non-vaccinated mice (Figure 3C-D; Supplemental Figure 5A-B). The increased numbers were much greater in pups born to dams who received the higher dose of vaccine (Figure 3B and D). In a separate experiment, significantly more neurons in the dentate gyrus of the vaccine-recipient offspring exhibited double-labeling for BrdU and NeuN compared to the matched controls, indicating long term survival of the newly generated neurons (Figure 3E-F). No significant differences were observed between pups of the vaccination or control groups in numbers of newly born astrocytes (Supplemental Figure 5C-D). There was no evidence of enhanced hippocampal cell proliferation or neuronal differentiation in the vaccination group at age 2 months (Supplemental Figure 6 A-F). Taken together, our data have demonstrated that neural precursor cell proliferation and neuronal differentiation are enhanced in adolescent pups in the context of maternal SARS-CoV-2 vaccination.

**Fig. 3.**
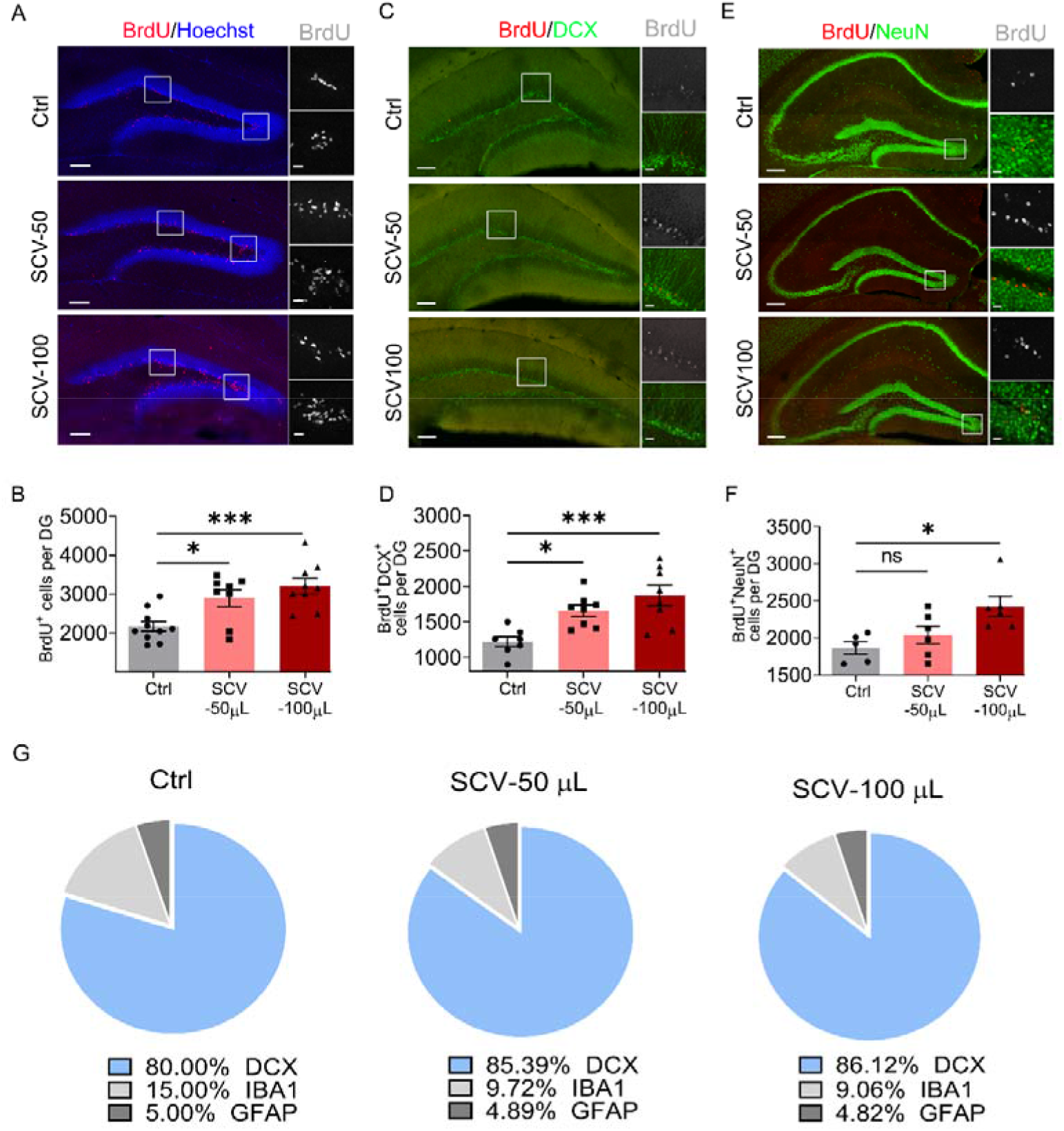
Maternal SARS-CoV-2 vaccination induces hippocampal cell proliferation and neuronal differentiation in pups at 1 month postnatally. (A) Representative confocal micrographs of the newborn cells, labeled by proliferative marker (BrdU) in the dentate gyrus (DG) of pups from SARS-Cov2-vaccinated pregnant mice with the inoculated doses of 50 (SCV-50) μL and 100 (SCV-100) μL and the vehicle controls. Two higher magnification (63 × oil immersion Len) images of the inset boxes in A were shown on the right (white). The image above was related to a granular layer, and the image below was related to a hilus. (B-C) Representative confocal images of the DG in each group showing BrdU (red) and DCX (green), BrdU (red) and NeuN (green). (D) The number of BrdU^+^ cells were analyzed in the unilateral DG of pups 24 h after the first BrdU injection. Summary data are presented as mean ± s.e.m. \**p* < 0.05, *\*\*\*p* < 0.001, one-way ANOVA followed by Tukey HSD test; ns: non-significant; *n*= 8-10 pups/group. Scale bar: 200 μm in A and B, 20 μm in higher images. (E-F) Quantification of the numbers of BrdU^+^/DCX^+^ and BrdU^+^/NeuN^+^ cells in the DG. The inset boxes in A and B were magnified (63 × oil immersion Len), as reflected by the higher images in the right (BrdU: white, NeuN: green). \**p* < 0.05 indicate significance by one-way ANOVA followed by the Tukey HSD test; *n*= 6-7 pups/group. Scale bar: 200 μm in A and B, 20 μm in higher images. (G) Percentage of BrdU^+^/DCX^+^, BrdU^+^/IBA1^+^ and BrdU^+^/GFAP^+^ cells to total BrdU^+^ in the DG among the three groups.

### Microglia are required for maternal SARS-Cov-2 vaccination-induced neuronal proliferation

We previously reported that microglia regulate developmental hippocampal neurogenesis in mice following maternal (influenza) or neonatal vaccination (hepatitis B)^16^ ^42^ ^52^. To investigate the potential effects of microglia on neuronal proliferation in the present study, 3-week-old pups from the SARS-Cov2-vaccinated dams were fed PLX chow (contains a pharmacologic inhibitor of receptors for CSF1, a growth factor critical for microglial survival). Microglial ablation was 95% compared with control microglial numbers (i.e., remaining Iba1-immunoreactive cells; Figure 4A-C). PLX chow did not affect astrocyte numbers (Figure 4D-E). Interestingly, microglia ablation abrogated the increase in hippocampal neurogenesis observed in pups of SARS-Cov2-vaccinated dams (Figure 4F-I). These data suggest that microglia mediate the pups’ neuronal proliferative response to maternal vaccination.

**Fig. 4.**
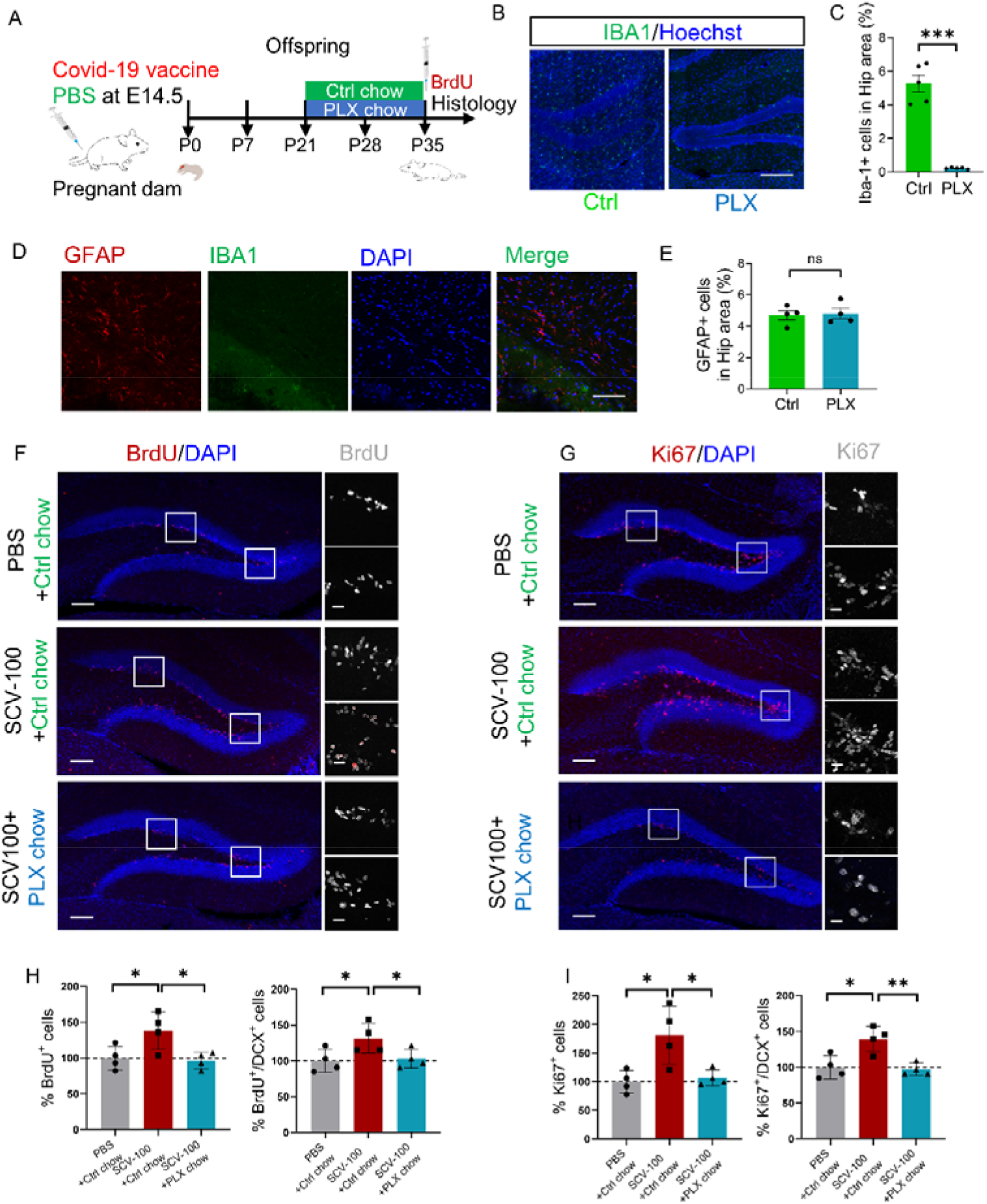
Microglia depletion abolishes maternal SARS-CoV-2 vaccination-induced hippocampal neurogenesis. (A) Schematic representation of the strategy to deplete microglia; (B, C) Representative images (*n* =4 sections per mice) and quantification of microglia marker IBA1 in the hippocampus from control (left) and PLX5563 chow-treated (right) mice. (D, E) Astrocytes survival was not affected in the brain slices from microglia-deleted mice. (F, G) Quantification of neuronal proliferation (BrdU^+^, Ki67^+^ in F, G) and differentiation (BrdU^+^ DCX^+^, Ki67^+^/DCX^+^ in F, G) in SCV pups and PBS pups after microglia deletion. Summary data are presented as mean ± s.e.m. \**p* < 0.05 indicate significance by one-way ANOVA followed by the Tukey HSD test; \**p* < 0.05, *\*\*p* < 0.01, *n*= 4-5 pups/group. Scale bar: 200 μm in B, F and G; 100 μm in D; 20 μm in higher images (right in F, G).

### Maternal SARS-CoV-2 vaccination increases microglia-proliferating neuron interaction

Next, we asked whether maternal vaccination activated microglia and how microglia affect hippocampal neuronal proliferation. As expected, the area of dentate gyrus occupied by microglia was significantly increased in the hippocampus of the vaccine offspring pups, relative to control pups (Figure 5A-D). Surprisingly, many more activated microglia surrounded and contacted proliferating cells in the dentate gyrus, where neuronal progenitors generate new cells during development and adulthood (Figure 5A-C and E). Numbers of microglia in the vaccinated and control pups did not differ significantly (Figure 5F). To further determine the microglial dependence of the observed enhancement in hippocampal neurogenesis, we evaluated the dentate gyrus of mice harboring genetically labelled microglia (Tmem119); dual positivity for BrdU and DCX identified proliferating neurons. 3D-reconstruction enabled visualization of direct contact between microglia and proliferating progenitor neurons, which was significantly more frequent in hippocampi of pups from vaccinated dams (Figure 5G). We classified the microglia-progenitor neuron interactions as follows: Type 1, two progenitor neurons (BrdU^+^DCX^+^) separated by a microglial process; Type 2, one progenitor neuron and one apoptotic cell (karyopyknosis) separated by a microglial process; Type 3, two progenitor neurons separated by a microglial soma (Figure 5H). The percentage of Type 1 interactions was greater in pups from vaccinated dams than in control pups (Figure 5I). Additionally, the area of dentate gyrus occupied by microglia positively associated with higher numbers of proliferating cells and immature newly generated neurons (Figure 5J). Because almost all microglia that were in contact with proliferating cells colocalized with immature neurons, our data suggested that maternal vaccination facilitated neuronal cell proliferation in the dentate gyrus, partially by augmenting interaction between microglia and proliferating neurons.

**Fig. 5.**
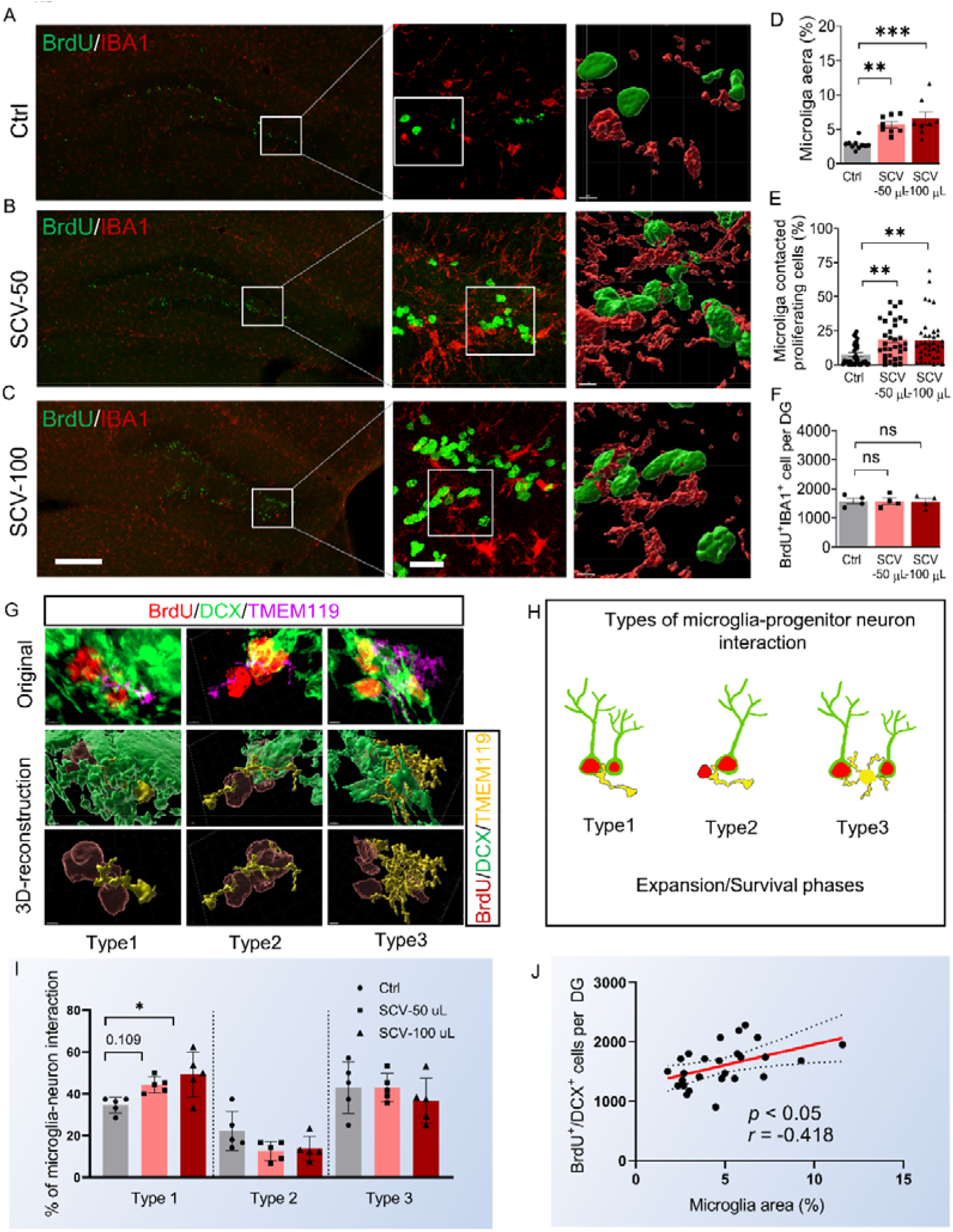
Maternal SARS-CoV-2 vaccination increases the microglia’s interaction with proliferating neurons. (A-C) Representative confocal images of the dentate gyrus showing BrdU (green) and IBA1 (red) in SCV pups and the vehicle controls. The higher magnification (63 × oil immersion Len) images of the left inset boxes in A-C were shown in the middle; 3D surface rendering reconstructions showing increased microglia (IBA1^+^) contacting proliferating cell (BrdU^+^) in right inset boxes using Imaris. (D-E) Quantification of microglia area and the percentage of microglia contacted with proliferating cells (BrdU^+^) in dentate gyrus in mice from A-C. (G) 3D surface rendering reconstructions of Tmem119^+^ microglia contacting BrdU^+^/DCX^+^ proliferating neurons using Imaris. Purple: Tmem119^+^ microglia (Original); Yellow: contacted microglia (3D-reconstruction) Green: DCX^+^ neurons; Red: BrdU^+^ proliferating cells. (H) Types of interactions between microglia and proliferating neurons during the expansion and survival phases of neuronal progenitors in development. (I) Percentage of microglia-proliferating neurons in different types of interactions in SCV-treated pups and PBS-pups (*n*=5 mice). (J) Positive correlation between the microglia area and the newborn neurons to in the DG in mice from A-C. The Spearman’s correlation analysis produces the P-values and correlation coefficients (R^2^). *n*= 8-12 pups/group in D; *n*= 33-38 cells/group in E. Scale bar: 200 μm in A-C; 3-15 μm in G (Original); 2-5 μm in G (3D-reconstruction). \**p* < 0.05 indicate significance by one-way ANOVA followed by the Tukey HSD test. \**p* < 0.05, \*\**p* < 0.01, \*\*\**p* < 0.001. ns. non-significant.

### *Cx3cr1* deficiency impairs SARS-CoV-2 vaccination-mediated microglia-proliferating neuron interaction

To further investigate immune mediators potentially implicated in microglia-proliferating neuron interactions, we quantified levels of 34 cytokines/chemokines in serum and hippocampal and cerebral cortex tissues. Unexpectedly, we found at 1 month after birth that pups of vaccinated dams had significantly less CXCL10 production and significantly more CCL22 production in comparison with pups of control dams (Supplemental Figure 7A-B). In the same sets of mice, the pups from vaccinated dams had, relative to the control pups, significantly higher hippocampal concentrations of CX3CL1, a chemokine ligand associated with enhancement of neurogenesis (Figure 6E). In contrast, hippocampal and cortical concentrations of the inflammatory chemokines CCL7 and CXCL16 (involved in recruitment and trafficking of leukocytes) were significantly lower in pups from vaccinated dams than in control pups (Supplemental Figure 7D-E). Interestingly, levels of most of the detected cytokines/chemokines (TNF-α, IL-1β, CCL11, CCL19, CCL22, CCL27, and CXCL11) were lower in the serum and hippocampus of pups from vaccinated dams than in control pups (Figure 6A-B; fold changes of SCV to Ctrl < 1). These findings suggest that SARS-CoV-2 vaccination during pregnancy causes long-term changes in prevailing cytokine and chemokine concentrations in the periphery and in brain, potentially creating a microenvironment conducive to neuronal proliferation and differentiation.

**Fig. 6.**
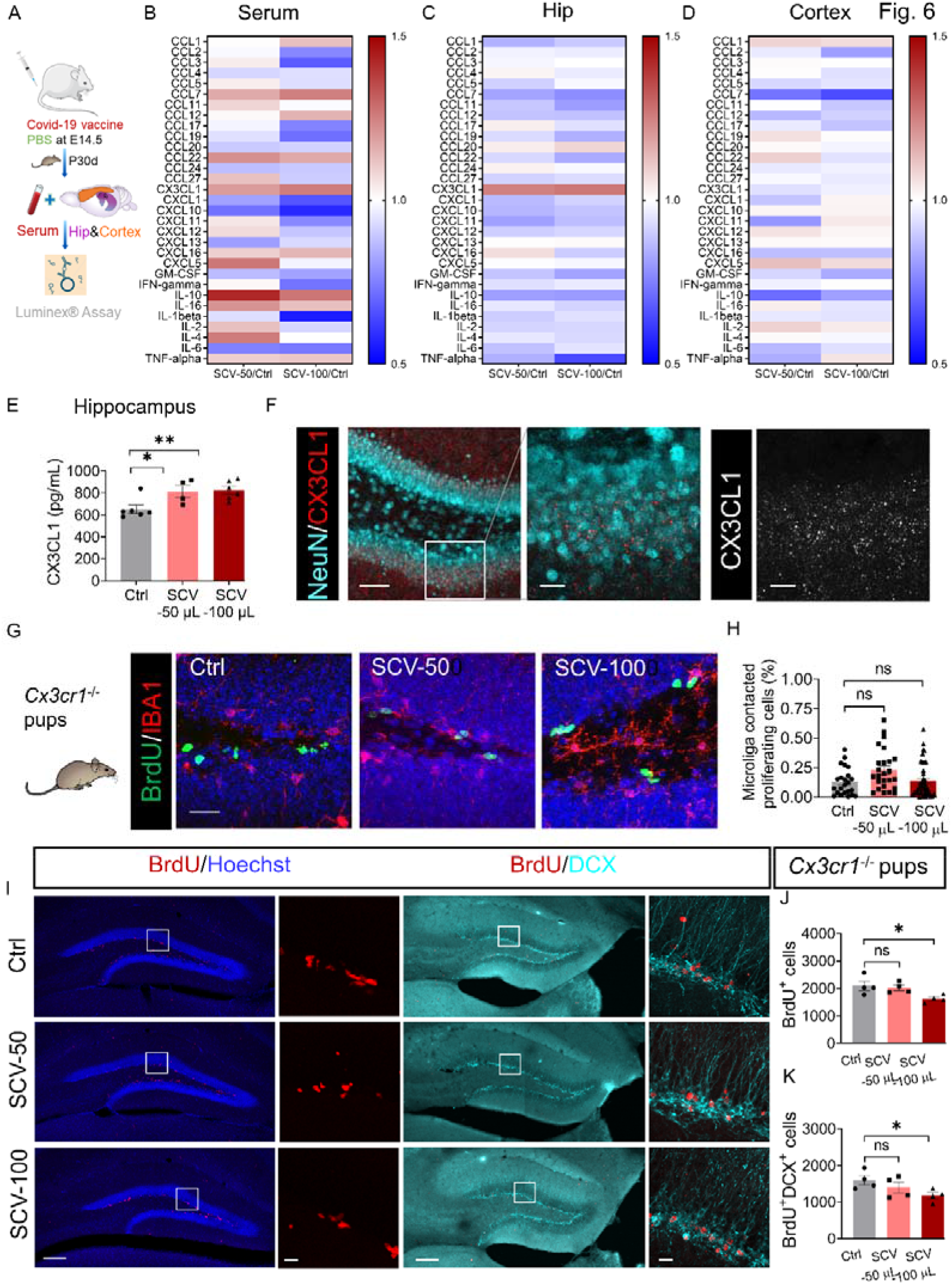
Maternal SARS-CoV-2 vaccination regulates chemokine/cytokine expression in the peripheral blood and the brain of pups. (A) Timeline for SARS-CoV-2 vaccination and Luminex Chemokine analysis. (B-D) The mean concentrations of 31 chemokine/cytokines were measured using the Bio-Plex Pro Mouse Chemokine Panel in the peripheral blood (B), hippocampal homogenate (C), and cortex homogenate (D) of pups (at age of 1 month) from SCV-50 dams (SCV-50), SCV-100 dams (SCV-100), and from the control dams (Ctrl). (E) CX3CL1 protein level was shown in hippocampus in pups from B-D. (F) Representative images of CX3CL1 (red) and NeuN (blue) in the DG. The inset box was magnified in F (middle); the CX3CL1 channel was shown in F (right). Scale bars: 100 μm (left); 50 μm (middle and right) in F. (G) *Cx3cr1* deficiency dampens SCV-mediated microglia-BrdU^+^ proliferating cell interaction. Scale bars: 50 μm in G (H) Quantification of microglia- BrdU^+^ cell interaction in the hippocampus of pups from SCV dams and the controls. (I) Representative images of proliferating cells (BrdU^+^, red, left) and newborn neurons (BrdU^+^DCX^+^, right) in pups from E. The higher magnification (63 × oil immersion Len) images of the left inset boxes in I were shown in right. Quantification of BrdU^+^ cells (J) and BrdU^+^DCX^+^ newborn neurons (K) in pups from I. Scale bars: 200 μm (left) and 50 μm (insert boxes) in I. Quantification of BrdU^+^ cells (I) and BrdU^+^DCX^+^ cells (J) in pups related to G and H. \**p* < 0.05 indicate significance by one-way ANOVA followed by the Tukey HSD test. \**p* < 0.05, \*\**p* < 0.01, ns. non-significant.

CX3CL1 signaling in neurons has been implicated in the promotion of adult neurogenesis in both dentate gyrus and subventricular zones ^23^ ^24^. By binding to its receptor CX3CR1, which is exclusively expressed on microglia, CX3CL1 regulates microglial motility in the developing central nervous system^53^ ^54^.

To determine whether CX3CL1 signaling in neurons is required for the maternal vaccination-mediated hippocampal neurogenesis and microglia-neuron interaction, we first confirmed that CX3CL1 protein was expressed in granular neurons of the pups’ dentate gyrus (Figure 6F). Next, we took advantage of the *Cx3cr1* deficiency in homozygous CX3CR1^GFP/GFP^ transgenic mice, which show no obvious differences in dentate gyral cell proliferation, relative to the *Cx3cr1* wild-type control mice (Supplemental Figure 8A-E). We found that hippocampi of pups born to *Cx3cr1* deficient mice that had received vaccine during pregnancy lacked evidence of interaction between microglia and proliferating neurons and had a low level of neurogenesis comparable to pups of non-vaccinated *Cx3cr1* deficient dams (Figure 6G-H). Thus, hippocampal neural cell proliferation and the emergence of immature neurons were impaired in offspring of *Cx3cr1* deficient mice receiving vaccination during pregnancy (Figure 6 G-H). Offspring of *Cx3cr1* deficient dams vaccinated with a high dose showed reduced proliferating cells and emergent immature neurons compared to control pups from non-vaccinated wild-type dams (Figure 6H).

Behaviorally, low-dose maternal SCV vaccination had no significant effects on locomotion and short-term memory functions in *Cx3cr1* deficient pups, but high-dose vaccination impaired these functions (Supplemental Figure 9A-B). Together, these data suggested that *Cx3cr1* deficiency hampers SCV vaccination-mediated interaction of microglia and proliferating neurons.

### Microglial *IFN-***γ***R1* deficiency impairs maternal SARS-CoV-2 vaccination-induced hippocampal neurogenesis

It has been reported that serum IFN-γ levels and numbers of IFN-γ^+^ SarsCov2-specific T cells are robustly elevated in healthcare workers who receive inactivated vaccine^38^ ^55^. To further investigate the possible molecular basis of maternal vaccination-induced hippocampal neurogenesis in young offspring mice, we measured the dams’ serum levels of IFN-γ protein at various time points after vaccination. There was a transient but significant increase at day 2 after vaccination compared to control dams (Figure 7A). The increase in IFN-γ was most obvious in the hippocampus of dams at day 2 and day 3 post-vaccination (Figure 7B). Given that a specific microglia/macrophage subset in the brain of both humans and rodents highly expresses IFN-γ receptor 1 (Figure 7B-C), we hypothesized that the impact of maternal vaccination on I month postnatal hippocampal neurogenesis and working memory is mediated by microglia through IFN-γ signaling. On investigating the effect of maternal SARS-CoV-2 vaccination on offspring of mice with conditional knockout of the *IFN-*γ*R1* gene in microglia, we found that maternal immune activation-mediated microglia activation (Figure 7E-F) and neuronal proliferation and differentiation in the postnatal hippocampus (Figure 7G-J) was abrogated. Furthermore, pups from vaccinated dams with selective ablation of microglial *IFN-*γ*R1* did not exhibit increased exploratory behavior toward novel objects or arms in the NOR task and Y maze task (Figure 7K-L). Together, these data suggest that the absence of IFN-γ signaling to microglia prevents the transiently increased short-term memory function observed in 1 month old postnatal pups of dams receiving vaccination during pregnancy.

**Fig. 7.**
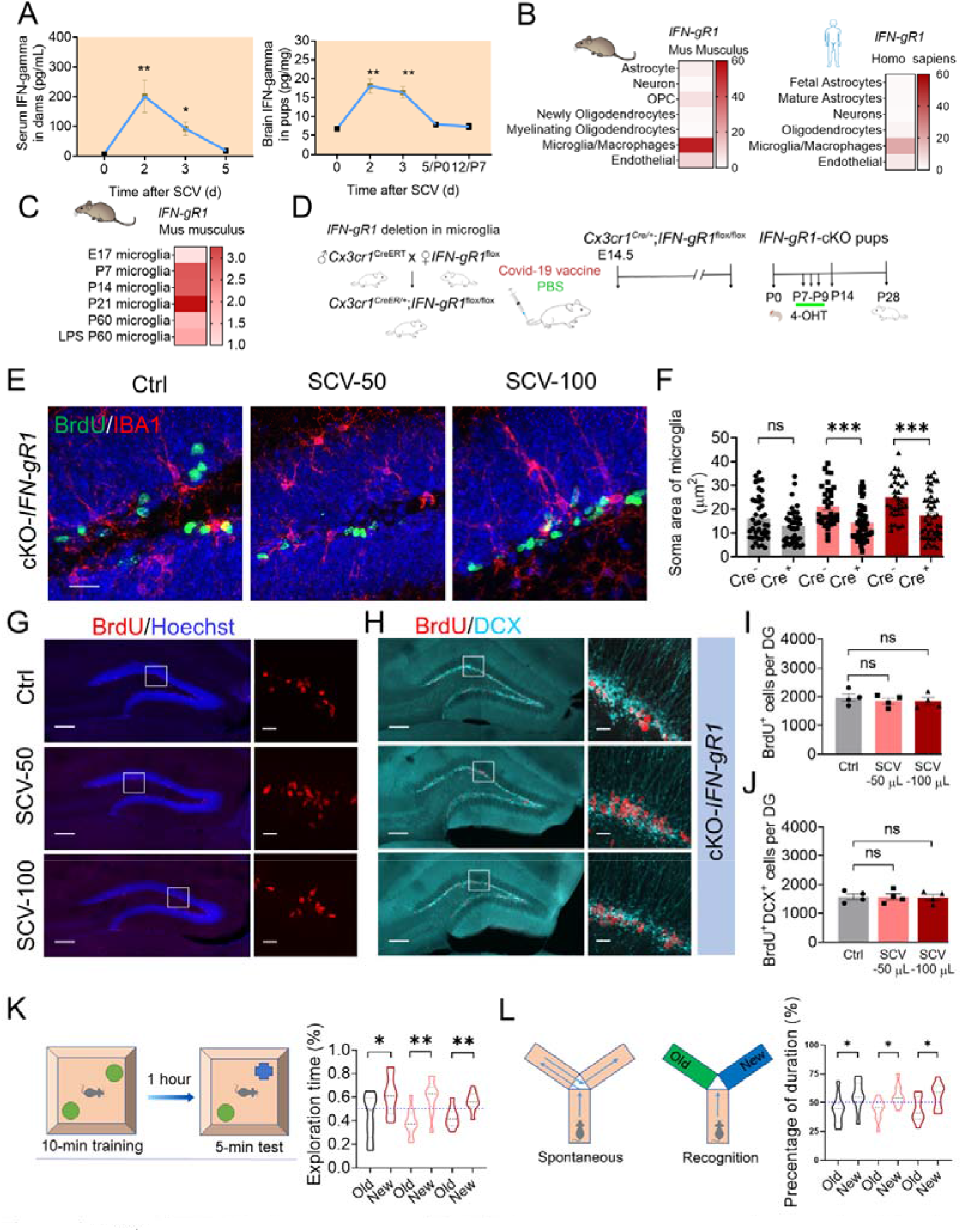
Absence of IFN-gR1 dampens maternal SARS-CoV2 vaccination-induced neuronal proliferating and differentiation. (A) *IFN-gR1* transcription in different cell populations in CNS of rodent and human being. (B) Quantification of IFN-γ protein levels in peripheral blood in dams after SARS-Cov-2 vaccination (left) and their pup’s brain (right). (C) *IFN-gR1* transcription in microglia/macrophages during development in rodent. (D) Schematic representation of the strategy to deplete *IFN-gR1* in CNS-resident microglia of pups, PBS, or SARS-CoV-2 vaccine (50 μL,100 μL, i.m.) administration at E14.5, and 4-OHT administration at P7 to P9 (1 mg/kg, 3 times i.p injection). Representative images of microglia contacted proliferating cells (E) and quantification (F) in controls and SCV pups after conditional *IFN-gR1* knockout. Scale bars: 50 μm in E. (G) Representative images of proliferating cells (BrdU^+^, red in G) and newborn neurons (BrdU^+^DCX^+^ in H) in pups from E. Scale bars: 200 μm (left) and 50 μm (insert boxes) in G and H. Quantification of BrdU^+^ cells (I) and BrdU^+^DCX^+^ cells (J) in pups related to G and H. NOR task (K) and Y maze test (L) for short-memory assessment were performed in pups from SCV dams and the controls after *IFN-gR1*deficentiy in microglia. New: novel object. Old: old object in K; New: novel arm. Old: old arm in L. one-way ANOVA followed by the Tukey HSD test in I and J; Student’s *t* test in A, F, K and L; \**p* < 0.05, \*\**p* < 0.01, \*\*\**p* < 0.001. ns. non-significant.

### Transcriptional signatures of BNT162b2 vaccination, another COVID-19 vaccine used in humans

To explain the possible derivation of IFN-γ-producing cells in the periphery after SARS-CoV-2 vaccination, we re-analyzed a single-cell sequencing database originally presented by Arunachalam et al.^47^, based on BNT162b2 vaccination of human subjects. We examined 242,202 cells from the database and segregated into 18 cell clusters (Figure 8A). Across several cell clusters, there was a notable range of differentially expressed genes (DEGs), with a 5 to 10-fold increase in expression observed on post-vaccination day 1 compared to day 0 (pre-vaccination). This contrasted with the gene expression on days 2 and 7, indicating that the most significant changes in gene expression occurred shortly after vaccination (Figure 8B). Gene-set enrichment analysis (GSEA) revealed that both BNT162b2 vaccination induced IFN-γ related module with time in the selected 8 cell clusters (Figure 8C), the curves of the IFN-gamma-related module score is consistent with the increased serum IFN-γ levels detected by ELISA (Figure 8A, below). In addition, IFN-γ signaling pathway gene expression, *IRF1* and *STAT1* were significantly increased in cDC2 cluster, CD14 monocyte, CD16 monocyte clusters and CD4 T cluster, CD8 T cluster, cDC1 cluster at day 1 post-vaccination as compared with day 0, respectively (Figure 8D-G). GSEA also revealed, at day 1 post-vaccination compared with pre-vaccination, enhancement of antibacterial humoral response in B cell cluster, antigen processing via MHCI and cellular response to type II interferon in CD8 T cluster, antigen presentation via MHCII, T cell cytokine production and T cell-mediated immunity in CD14 monocytes. Most noteworthy were substantial enrichment of modules associated with Type II interferon, T cell immunity and antigen presentation when genes were ranked based on their correlation with IFN-γ on day 1 (Figure 8H). When the CD8 T cell cluster was selected for analysis, the top-ranked pathways included lymphocyte-mediated immunity, positive regulation of cytokine production, lymphocyte migration, and response to type II interferon (Figure 8I). These published human post- BNT162b2 vaccination data suggest a pivotal role for IFN-γ in promoting antigen presentation by monocytes, enhancing T cell immunity, and boosting the humoral antibacterial response in B cells following administration of this vaccine.

**Fig. 8.**
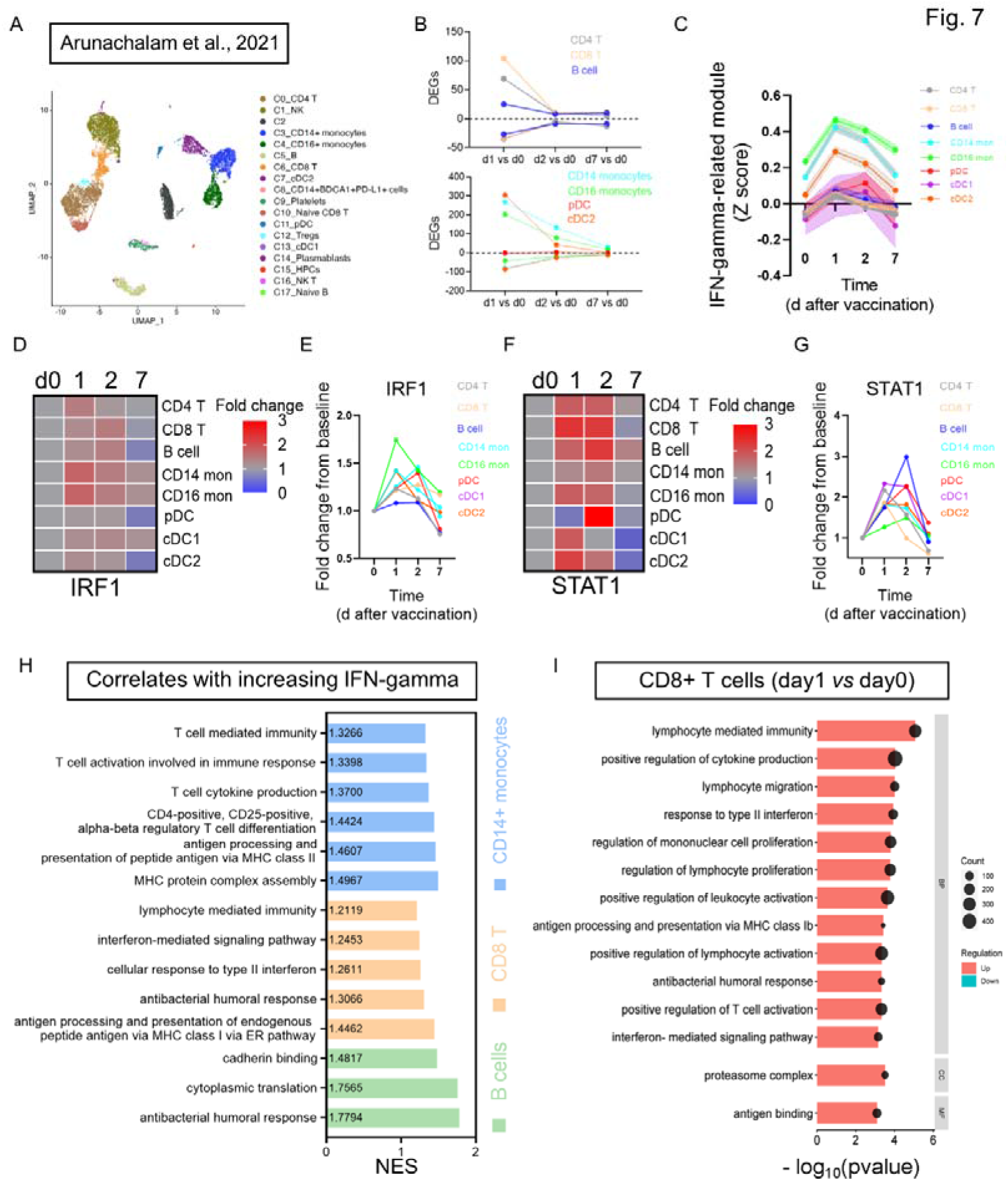
Single cell RNA-sequence data of BNT162b2 vaccination in human. (A) UMAP representation of PBMCs subtypes identified by single-cell transcriptional profiling from Arunachalam et al.’s paper. (B) Number of differentially expressed genes (DEGs) (absolute log_2_FC > 0.1 and adjust *P* < 0.05) at day 1, day 2 and day7, compared with day 0. (C) IFN-gamma-related module score before vaccination (day 0) and after vaccination (day 1, day 2 and day7) based on the Interferon-mediated Signaling Pathway (GO:0140888). (D, E) Heatmap for interferon regulatory factor 1 (*IRF1*) gene in cell types at each time point before and after SARS-CoV-2 vaccination. (F, G) Heatmap for *STAT1* gene in cell types at each time point. (H) Gene-set enrichment analysis (GSEA) results of the significantly upregulated immune-related pathways in B cells, CD8 T cells and CD14 monocytes at day 1/day2 compared to day 0. (I) Lymphocyte- and type LJ interferon-mediated pathways were upregulated in the CD8 T cells between days 1 and day 0 by GSEA analysis.

## Discussion

The data we present show that administration of inactivated SARS-CoV-2 vaccine to pregnant mice leads to an increase in hippocampal neurogenesis and enhancement of working memory in pups at age 1 month but not at 2 months. As in previous studies, this study confirmed that maternal vaccination had no adverse effect on the physical condition of dams or their pups^13^ ^14^. Of note, SARS-CoV-2 IgG was detected in serum of 1-month-old pups born to dams receiving inactivated SARS-CoV-2 vaccine during the early third trimester. Experiments employing mice with conditional knockout of microglial *IFN-rR1* or general knockout of *Cx3cr1* genes demonstrated a role for microglia-neuron interaction in mediating the transient hippocampal neurogenesis and memory enhancement in off-spring of mice receiving SARS-CoV-2 vaccine in late pregnancy.

### The safety and effectiveness of SARS-CoV-2 vaccination during pregnancy

Many pregnant or breastfeeding women are hesitant to accept SARS-CoV-2 vaccine due to concerns about adverse events, including fertility or spontaneous abortion ^56^ ^57^. The data for our mice indicated no adverse effect of maternal vaccination on the number of embryos, birth weight, or their growth and development. These data are consistent with a recent study of a human vaccination trial using another inactivated SARS-CoV-2 vaccine (CoronaVac, Sinovac Life Sciences)^13^. Of translational importance, the current study has documented that a maternal protective IgG response is established by vaccination with SARS-CoV-2 RBD protein in late pregnancy and is detectable in offspring at 1 month of age^58^. No abnormal behavior (anxiety and social activities as evidenced by OFT and three-chamber social tests) was observed in pups whose mothers had received the vaccine. Further supporting the safety profile of SARS-CoV-2 vaccination during pregnancy, levels of most of the measured cytokines/chemokines remained low in both serum and hippocampus of the (SARS-CoV-2 vaccinated) pups. Our findings add to the growing body of evidence supporting the safety and efficacy of SARS-CoV-2 vaccination during pregnancy.

### Maternal SARS-CoV-2 antibodies are transferred across the placenta

Previous research has demonstrated that maternal antibodies specific for influenza or SARS-CoV-2 are transferred to progeny across the placenta and postnatally through lactation^59^ ^60^. Maternal protective antibody is thought to reduce the risk of influenza in the first months of an infant’s life^61–63^. Maternal SARS-CoV-2 vaccination is anticipated to provide beneficial protection by transferring IgG to newborns^58^. However, in human pregnancy studies, higher titers of SARS-CoV-2 antibodies were detected in cord blood of neonates whose mothers received COVID-19 vaccine (mRNA or inactivated virus protein vaccine) but not during lactation^64^. Not surprisingly, in the present study we detected higher titers of SARS-CoV-2-specific antibodies in the serum of vaccinated dams compared with their pups. The SARS-CoV-2 RBD-IgG inhibition ratio of the dams was 2-fold higher than that of their progeny. Recent data have suggested that transfer of SARS-CoV-2-specific IgG is compromised in the third trimester of pregnancy, and have implicated perturbance of Fc glycosylation, a critical determinant of maternal-fetal IgG transfer ^59^. Therefore, this sharp reduction in the titers of pup IgG questions the appropriate timing of SARS-CoV-2 vaccination during pregnancy^65^. Future studies need to assess whether maternal vaccination during early or later pregnancy would yield a higher titer of SARS-CoV-2 IgG in pups.

### Maternal SARS-CoV-2 vaccination increased hippocampal neurogenesis and working memory in pups

The SARS-CoV-2 virus is known to invade the central nervous system (CNS) and cause neurological and neuropsychiatric complications ^66–69^. Patients who have experienced COVID-19 infection are more likely to have impaired neurogenesis ^69–71^ and cognitive decline ^72^ ^73^. However, these data were not focused on pregnant women, which is the focus of our study. Here, we have shown that SARS-CoV-2 vaccination in the second half of pregnancy not only transiently promoted neuronal proliferation and neuronal differentiation in postnatal pups, but also improved their working memory faculties. We noted that the postnatal enhancement of hippocampal neurogenesis by maternal vaccination was only observed during the adolescent period and did not extend into adulthood. This trend is consistent with our earlier published observations on the effects of influenza vaccination during early pregnancy^15^ and of live Bacille Calmette-Guerin (BCG) vaccination in neonates ^74^. Our present study provides compelling evidence for neuro-immune interactions, which has been well established over the past two decades ^75^ ^76^.

### Microglia-neuron interactions drive neurogenesis during brain development

Microglia-neuron crosstalk has been extensively studied in the course of physiological synaptogenesis^77–79^ and neurodegenerative synaptopathies^80–82^. In adulthood, microglia can shape hippocampal neurogenesis through phagocytosis of apoptotic newborn cells ^83^. Less is known about how microglia regulate proliferating neurons in during hippocampal DG development. Our present study has revealed the importance of CX3CL1/CX3CR1 signaling for microglial chemotaxis and neuronal proliferation during brain development. It has been reported that the intracellular domain of CX3CL1 enhances adult neurogenesis and replenishes lost neurons in mouse models of neurodegenerative diseases^23^. *Cx3cr1* deficiency impairs synaptic plasticity and hippocampal cognitive function in adulthood (at age 3 months)^84^. Our findings emphasize the role of CX3CL1 and its receptor in microglial-neuron interactions mediated by chemotaxis, thereby contributing to neurogenesis via IFN-γ-dependent mechanisms during brain development. Our findings suggest that cerebral IFN-γ derived from a peripheral source, targets microglial receptors to induce microglial activation. Future studies are needed to determine whether IFN-γ/IFN-γR1 couples with neuronal-derived CX3CL1/CX3CR1 to exert a chemotactic role in microglia-neuron interaction.

### A possible mechanism for maternal SARS-CoV-2 vaccination impacting neurogenesis

Our earlier studies have demonstrated that peripheral immune activation induced by BCG or influenza vaccination promotes hippocampal neurogenesis^16^ ^43^. We have proposed plausible mechanisms involving elevation of BDNF expression by neurons or astrocytes ^43^ and activation of microglia expressing IGF-1, another neurotrophic factor^15^. We have also observed that maternal influenza vaccination promotes IFN-γ-dependent upregulation of adhesion molecules and chemokine production in the choroid plexus (CP) of pups with recruitment of T lymphocytes and regional neuronal cell proliferation and transient enhancement of working memory^42^. These maternal vaccination-related benefits were eliminated if T lymphocyte in the brain were ablated by administering T cell receptor (CD3)-neutralizing IgG^42^. It is pertinent, in the human context, that expression of genes regulating lymphocyte migration, T cell differentiation and regulation of T cell proliferation pathway were enriched by GESA analysis of PBMC following maternal COVID-19 vaccination^47^. Therefore, it might be worthwhile to investigate SARS-CoV-2 T-cell mobilization in the brain barrier or cerebrospinal fluid after COVID-19 vaccination^85^, which enhances hippocampal neurogenesis in the present study. Whether maternal influenza vaccination and SARS-CoV-2 vaccination shared common mechanisms for pro-neurogenesis effects, including involvement with BDNF and IGF-1 signaling still need to be resolved. Based on our previous research, a recent review paper concluded that COVID-19 vaccination may enhance hippocampal neurogenesis ^17^. Here, our data provided behavioral and cellular evidence to validate the hypothesis in a pregnant mouse model.

## Limitations

This project primarily emphasizes animal studies and lacks clinical data based on human responses after COVID-19 vaccination. It’s important to acknowledge that transcriptional signatures of PBMCs, as revealed by single-cell RNA sequencing, in individuals vaccinated with the Pfizer-BioNTech mRNA vaccine (BNT162b2, against clone SARS-CoV-2 derived from 2019-nCoV/USA_WA1/2020 strain) may differ from those associated with the inactivated SARS-CoV-2 vaccine (BBIBP-CorV, Sinopharm, derived from SARS-CoV-2 strain 19nCoV-CDC-Tan-HB02) used in the present study.

Therefore, the inclusion of more clinical evidence is essential to substantiate the conclusions drawn in this study. In addition, this project also lacks the data of cell-derive of IFN-γ both in the periphery and the brain. However, although IFN-γ can be released by peripheral monocytes, DCs and T cells after COVID-19 vaccination, as reflected by single-cell RNA data from human’s PBMC.

## Conclusions

Our study highlights the safety and efficacy of inactivated SARS-CoV-2 vaccination during pregnancy. Additionally, our findings shed light on a fundamental mechanism through which adaptive immune responses influence neuronal development via the interaction of microglia and proliferating neurons. Microglial IFN-γ and CX3CR1 signaling are implicated in regulating this interaction.

## Acknowledgments

We thank all the members of Prof. Zhilin Yao’s lab for discussion, Ms. Tuo Zhou and Mr. Junjie Chen for technical support, the Core Lab Platform for Medical Science for technical assistance and support from the Animal Center of Zhongshan School of Medicine, Sun Yet-sen University and Prof. Vanda Lennon of Mayo Clinic for critically reviewing the manuscript.

## Authors’ contributions

Fangfang Qi, Zhibin Yao, Dandan Hu and Kaihua Guo contributed to the design and supervised this study and the analyze the data. Fangfang Qi, Jiaoling Tang and Juntao Zou wrote the manuscript. Lingxiao Wang performed the single cell sequencing analysis. Ashutosh Kumar contributed to valuable comments and discussion for this project. Fangfang Qi, Jiaoling Tang, and Hao Liu performed the experiments. Fangfang Qi wrote the paper.

