## Supplemental Material for "SARS-CoV-2 vaccination during pregnancy enhances hippocampal neurogenesis and working memory in offspring via IFN-gamma responsive microglia"

Fig. S1 Effects of maternal SARS-CoV-2 vaccination on litter size and physical development

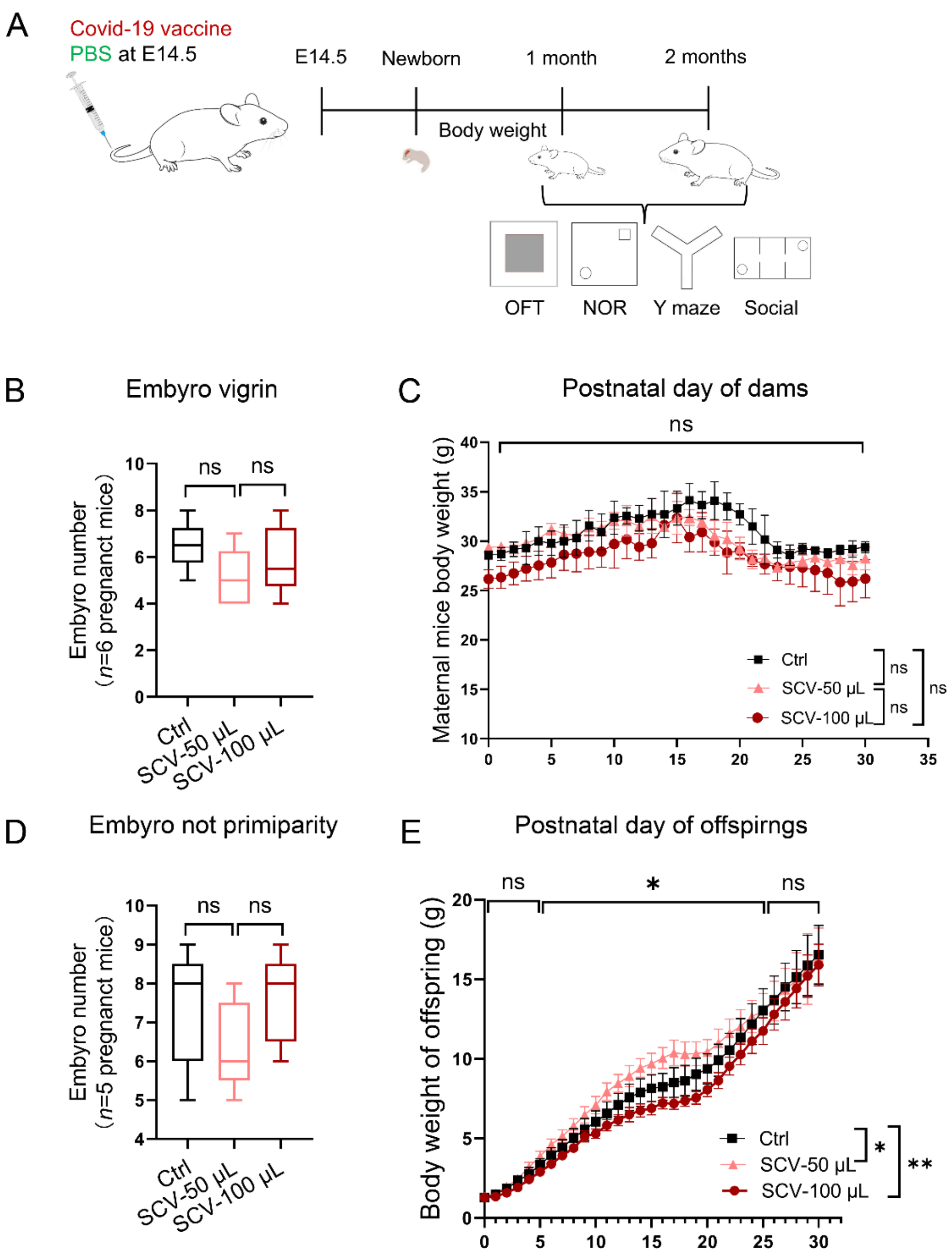


**Fig. S1 Effects of maternal SARS-CoV-2 vaccination on litter size and physical development**

(A) Timeline of maternal SARS-CoV-2 vaccination, body weight, and behavioral tests, including OFT, social preference, and NOR tests. Viable embryos were obtained from primiparous (B) or non-primiparous (D) pregnant mice. ns: non-significant, one-way ANOVA followed by the Tukey HSD test, *n* = 4-5 dams/group. (C) The body weights of

SARS-CoV-2-vaccinated pregnant mice and the controls were measured each day after delivery. (E) The body weight of the pups from SARS-CoV-2-vaccinated or PBS-treated dams was obtained each day after birth. Two-way repeated ANOVA analysis for body weight, *$p < 0.05$, $n$ =12-14 pups/group.

Fig. S2 Effects of maternal SARS-CoV-2 vaccination on offspring exploratory and short-term memory function at 1 month postnatally

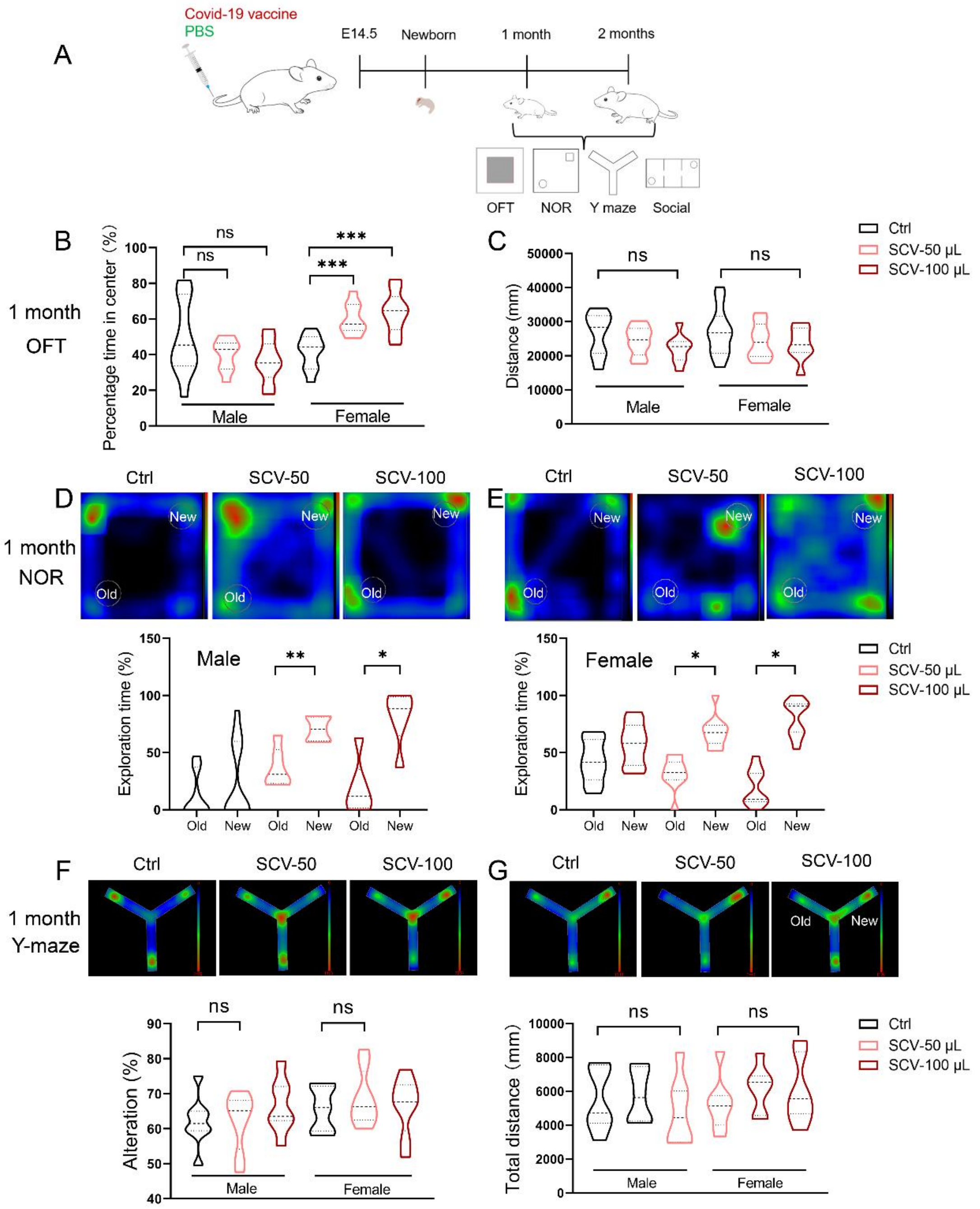


**Fig. S2 Effects of maternal SARS-CoV-2 vaccination on offspring exploratory**

**and short-term memory activities at 1 month postnatally**

(A) Experimental procedure: The exploratory behavior, object recognition memory ability, and social activity of pups from vaccinated dams and the controls were assessed in OFT and NOR tests at 1 month postnatally. Exploratory activities in the center area (B) Total distance (C) of male and female pups from SCV dams and the controls were conducted in OFT at 1 month postnatally. One-way ANOVA followed by Tukey HSD test, $n$= 4-5 dams/group. (D-E) Representative Heatmap of movement tack and quantification of male (D) and female (E) pups from SCV dams (two doses) and the controls in NOR test. New: novel object. Old: old object. (F and G) Representative heatmaps of male and female pups from SCV dams (two doses) and the controls in the Y-maze test. Quantifying spontaneous alteration (F) and total distance (G) in male and female pups during the 5-min duration of the Y-maze test. Paired Student's two-tailed *t-test* for the analysis in NOR test. One-way ANOVA followed by Tukey HSD test in OFT test and Y maze test; *$p < 0.05$, **$p < 0.01$; ns: non-significant.

Fig. S3 No effects of maternal SARS-CoV-2 vaccination on short-memory ability and social behavior in offspring 2 months after birth

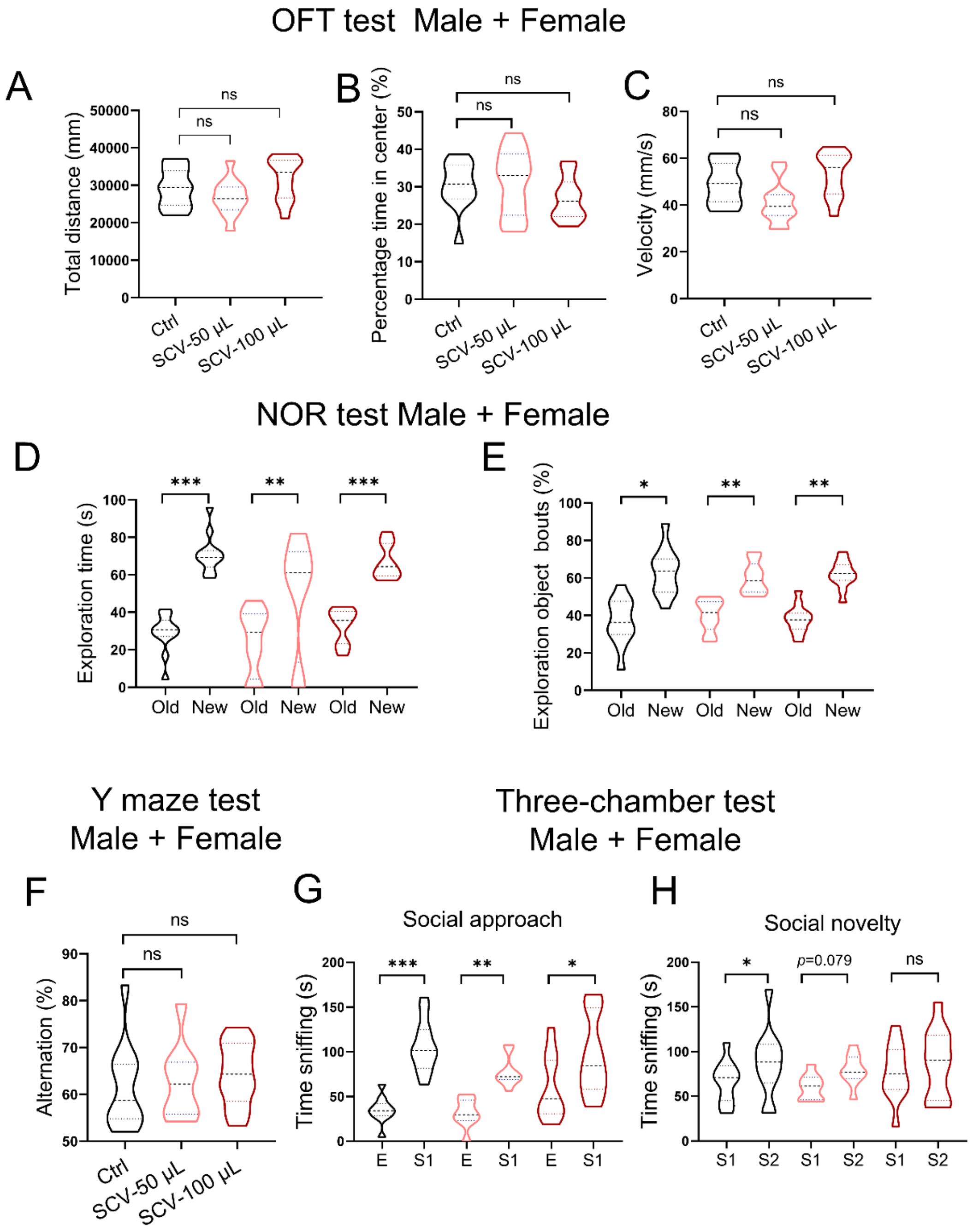


**Fig. S3 No effects of maternal SARS-CoV-2 vaccination on short-memory ability and social behavior in offspring 2 months after birth**

(A-C) The exploratory behavior, including total distance (A), percentage time in center (B), and velocity (C) of pups from SARS-CoV2 vaccinated dams and PBS-treated controls were measured in OFT at 2 months postnatally, *$p < 0.05$ indicate significance by one-way ANOVA followed by Tukey HSD test; ns: non-significant; $n$=12 pups/group.

(D-F) Novel object recognition memory of pups from SARS-CoV2 vaccinated dams and PBS-treated controls were assessed in the NOR test at the age of 2 months (D), regardless of sex factor. All pups performed normal recognition memory at 2h after training (E and F). *$p < 0.05$ indicates significance by paired Student's *t-test*. **$p < 0.01$, ***$p < 0.001$; $n$=9-12 pups/group in D-F. (G) Quantification of spontaneous alteration in pups from SCV dams and the controls during 5-min duration in Y-maze test, regardless of sex factor. (H) Duration spent in sniffing stranger and empty cage by 2-month pups in social approach test, regardless of sex factor. (I) Duration spent in sniffing stranger 1 and stranger 2 exploration time by 2 months, regardless of sex factor. Paired Student's two-tailed *t-test*. *$p < 0.05$, **$p < 0.01$, ***$p < 0.001$; ns: non-significant; $n$=9-12 pups/group. E: empty; S1: stranger 1; S2: stranger 2.

Fig. S4 The effects of maternal SARS-CoV-2 vaccination on offspring social behavior 1 month after birth

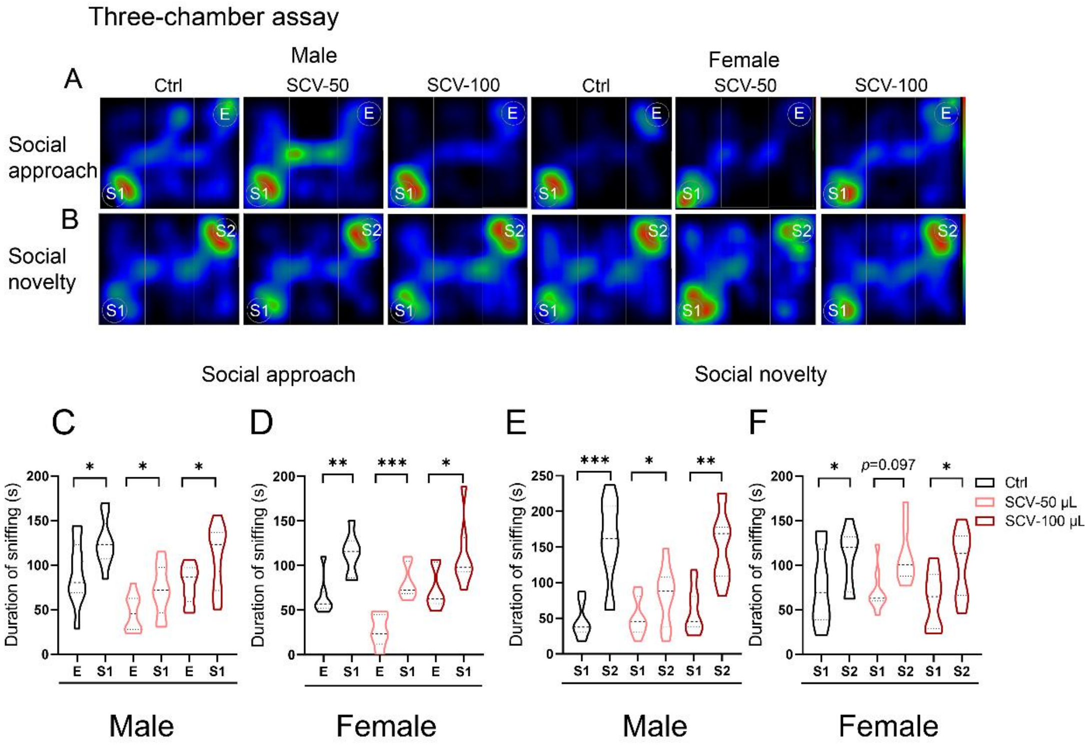


**Fig. S4 The effects of maternal SARS-CoV-2 vaccination on offspring social behavior 1 month after birth**

(A-B) Representative Heatmap of male and female pups during social approach (A) and social novelty (B) phases in the three-chamber assay. (C-D) Duration spent in sniffing stranger and empty cage by 1-month male and female pups in social approach test. (E-F) Exploration time spent sniffing stranger 1 and stranger 2 by 1-month male and female pups. The paired student's two-tailed *t-test*. *$p < 0.05$, **$p < 0.01$, ***$p < 0.001$; ns: non-significant. E: empty; S1: stranger 1; S2: stranger 2.

Fig. S5 Maternal SARS-CoV-2 vaccination differently affects hippocampal neuronal stem cell proliferation and astrocytes in offspring at 1 month postnatally

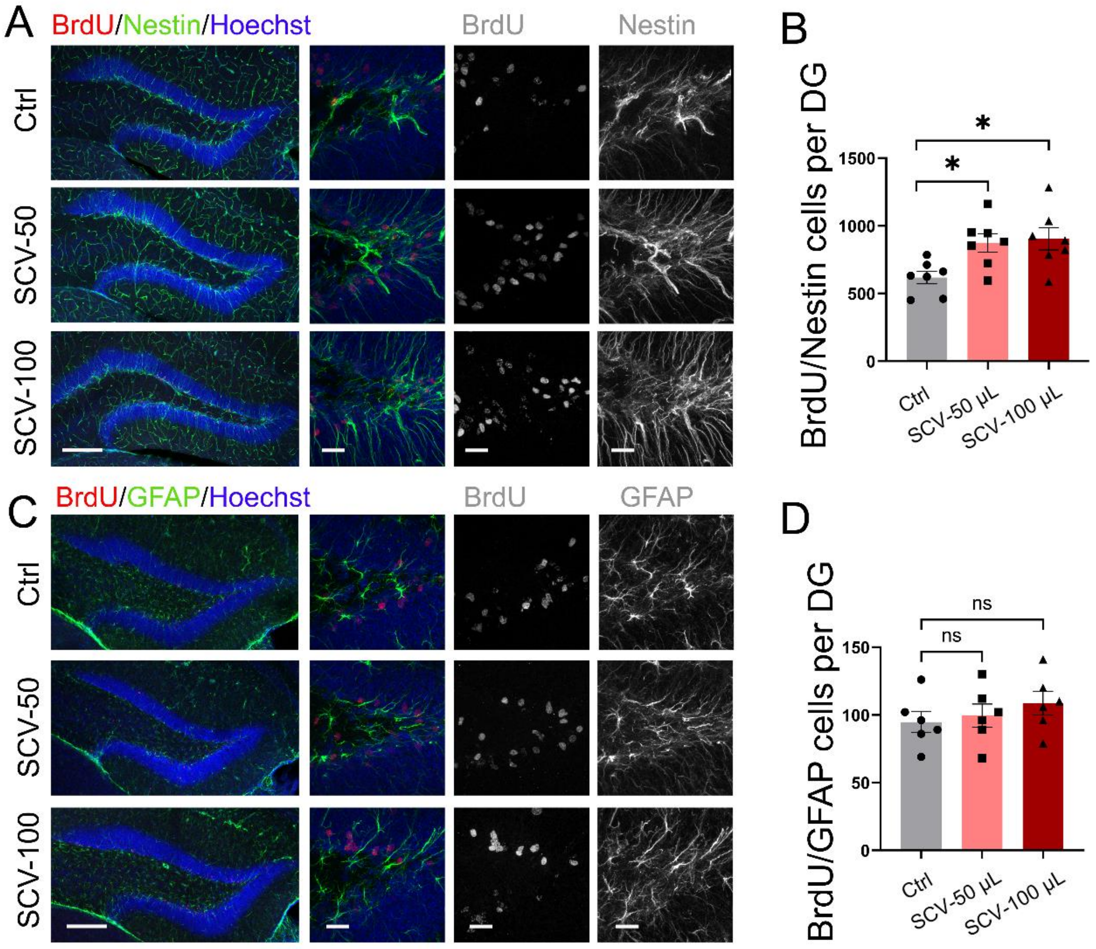


**Fig. S5 Maternal SARS-CoV-2 vaccination differently affects hippocampal neuronal stem cell proliferation and astrocytes in offspring at 1 month postnatally**

Representative images of immunolabelled newly neuronal stem cells (Brdu$^+$/Nestin$^+$ in A) and newborn astrocytes (BrdU$^+$/GFAP$^+$ in B) in DG of pups born to both SARS-CoV-2 vaccinated dams and PBS-treated control dams. (B and D) Quantification of the numbers of BrdU$^+$/Nestin$^+$ and BrdU$^+$/GFAP$^+$ cells in DG. *$p < 0.05$ indicates significance by one-way ANOVA followed by Tukey HSD test; ns: non-significant; $n$= 6-7 pups/group. Scale bar: 200 μm in A and C (left column, lower power lens), 20 μm in A and C (higher power lens).

Fig. S6 Maternal SARS-CoV-2 vaccination does not significantly alter hippocampal cell proliferation and differentiation in offspring at 2 months postnatally

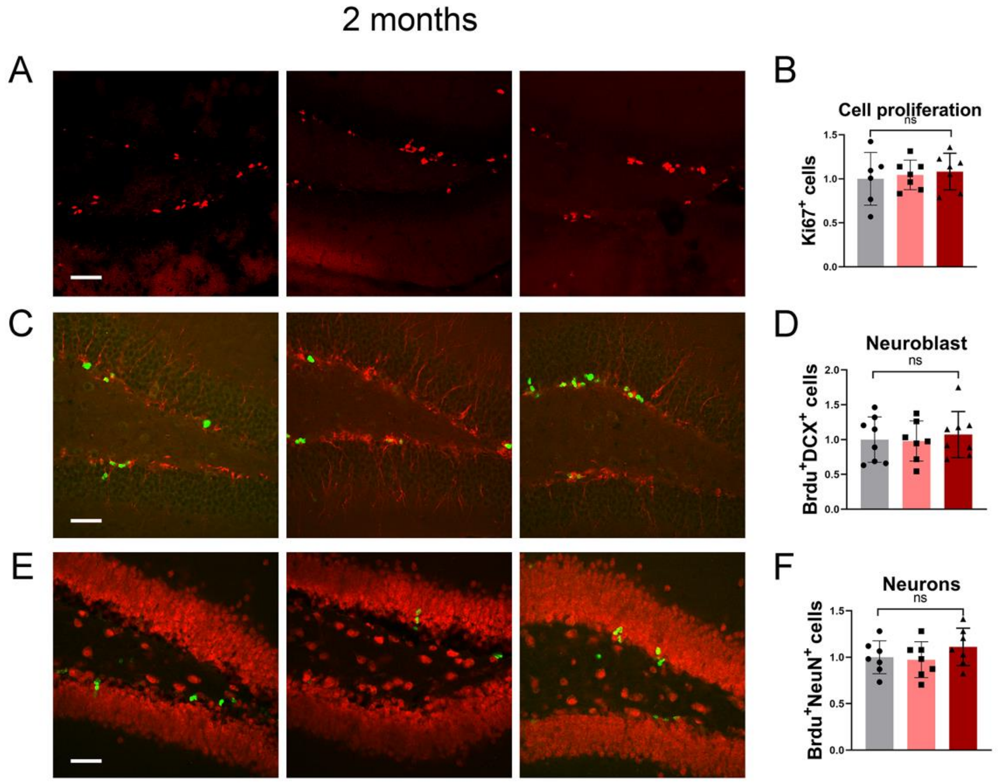


**Fig. S6 Maternal SARS-CoV-2 vaccination does not significantly alter hippocampal cell proliferation and differentiation in offspring at 2 months postnatally**

Representative images of immunolabelled Ki67$^+$ cells (A), BrdU$^+$/DCX$^+$ cells (C), BrdU$^+$/NeuN$^+$ cells (E) in DG of pups born to both SARS-CoV-2 vaccinated dams and PBS-treated control dams. Fold change in immunolabelled cells in the unilateral DG.

For Ki67⁺, BrdU⁺/DCX⁺ and BrdU⁺/NeuN⁺ cells, *$p < 0.05$ indicates significance by one-way ANOVA followed by Tukey HSD test; ns: non-significant; $n$= 7-8 pups/group. Scale bar: 50 μm in A, C and E.

Fig. S7 Maternal SARS-CoV-2 vaccination changes the expression of chemokine/cytokine in the serum and the brain of offspring at 1 month postnatally

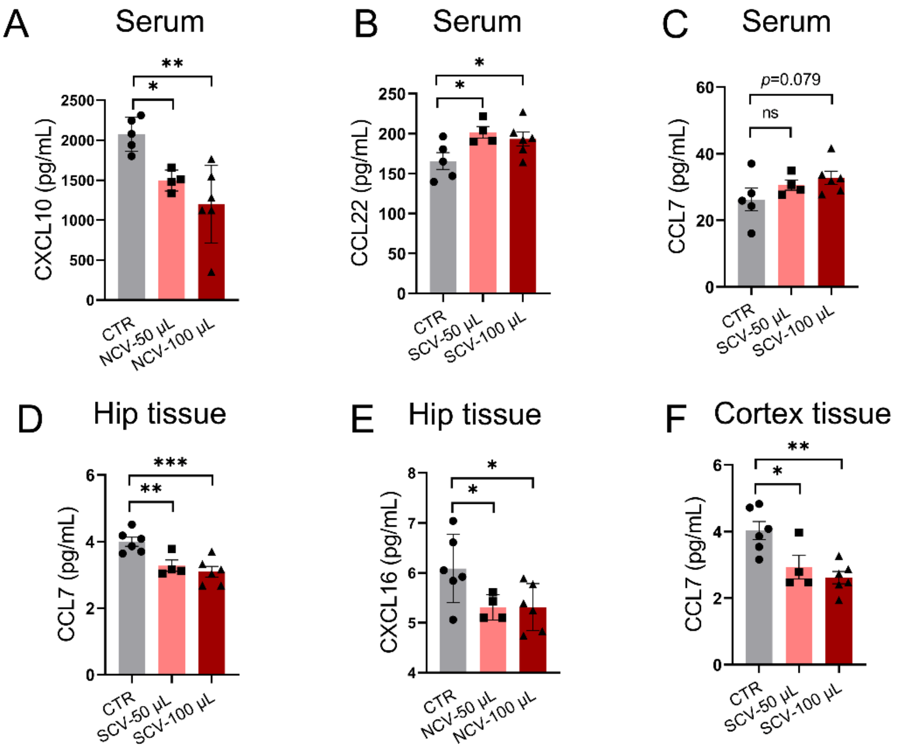


**Fig. S7 Maternal SARS-CoV-2 vaccination changes the expression of chemokine/cytokine in the serum and the brain of offspring at 1 month postnatally**

(A-C) The levels of cytokine CXCL10, CCL22, and CCL7 in the serum were shown by histograms, which showed significant differences from the pups born to SARS-CoV-2 vaccinated dams and PBS-treated controls. (D-F) The levels of cytokine CCL7 and CXCL16 in the hippocampus were shown in D and E. The levels of cytokine CCL7 in the cortex were shown in F, which were significantly changed in pups born to vaccinated dams, relative to the controls. *$p < 0.05$ indicates significance by one-way ANOVA followed by Tukey HSD test; *$p < 0.05$, **$p < 0.01$, ***$p < 0.001$; $n$ =6 pups in Control pups and SCV-100 pups; $n$=4 pups in SCV-50 pups.

Fig. S8 Absence of Cx3cr1 receptor does not affect hippocampal cell proliferation at 1 month postnatally

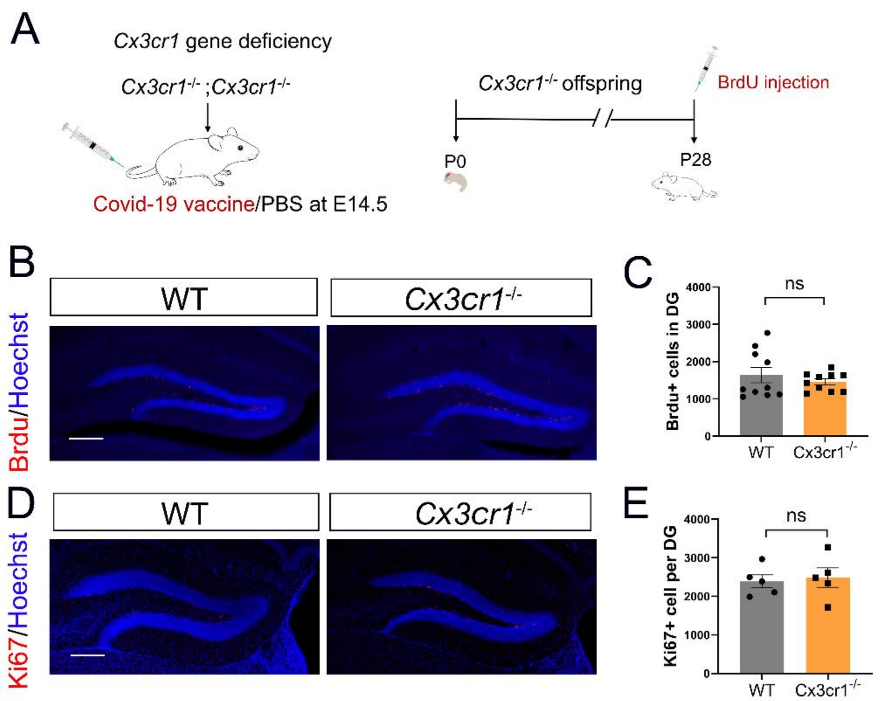


**Fig. S8 Absence of Cx3cr1 receptor does not affect hippocampal cell proliferation at 1 month postnatally**

(A) Schematic of *Cx3cr1* deficient pups' generation and the timeline of SARS-CoV-2 vaccination, BrdU injection (50 mg/kg, i.p.). (B) Representative images and (C) quantification of immunolabelled BrdU$^+$ cells in WT pups and *Cx3cr1* KO pups. (D) Representative images and quantification of Ki67$^+$ cells in mice from B and C. Scale bar: 200 μm in B and D. *$p < 0.05$ indicates significance by unpaired Student's two-tailed *t-test* for the analysis in C and E; ns: non-significant; $n$=10 pups/group in C, $n$=5 pups/group in E.

Fig. S9 Cx3cr1 deficiency impairs maternal SARS-CoV-2 vaccination-induced increase in short-memory function

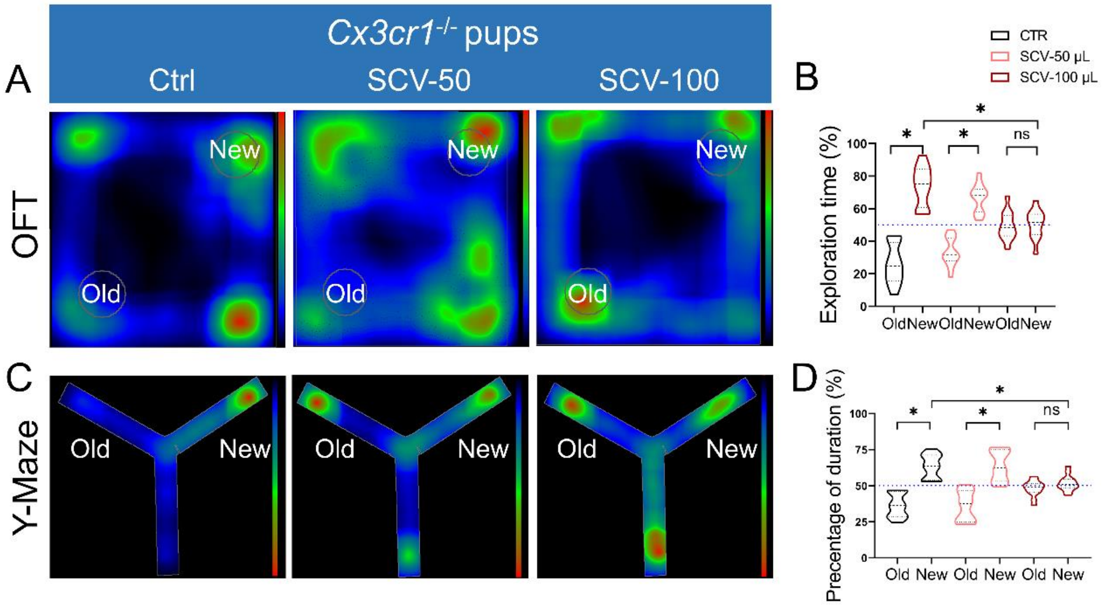


**Fig. S9 *Cx3cr1* deficiency impairs maternal SARS-CoV-2 vaccination-induced increase in short-memory function**

NOR task was conducted in the *CX3CR1* deficient pups from SARS-CoV-2 vaccinated dams and PBS-treated controls. (A) Representative Heatmap of movement tack and (B) quantification of preference to novel object and old object in NOR test. New: novel object; Old: old object. (C) Representative Heatmap of movement tack and (D) quantification of duration in the novel arm and old arm in the Y maze test. New: novel arm. Old: old arm. ns: non-significant; *$p < 0.05$ indicates significance by one-way ANOVA followed by Tukey HSD test; $n$=9-11 pups/group.
